# Increased sensitivity to strong perturbations in a whole-brain model of LSD

**DOI:** 10.1101/2021.01.05.425415

**Authors:** Beatrice M. Jobst, Selen Atasoy, Adrián Ponce-Alvarez, Ana Sanjuán, Leor Roseman, Mendel Kaelen, Robin Carhat-Harris, Morten L. Kringelbach, Gustavo Deco

**Affiliations:** Center for Brain and Cognition, Computational Neuroscience Group, Universitat Pompeu Fabra, Calle Ramón Trias Fargas 25-27, 08005 Barcelona, Spain; Department of Psychiatry, University of Oxford, Oxford, UK; Center of Music in the Brain (MIB), Clinical Medicine, Aarhus University, DK; Centre for Psychedelic Research, Department of Brain Sciences, Imperial College London, United Kingdom; Institució Catalana de la Recerca i Estudis Avançats (ICREA), Barcelona, Spain; Department of Neuropsychology, Max Planck Institute for Human Cognitive and Brain Sciences, Leipzig, Germany; School of Psychological Sciences, Monash University, Clayton, Melbourne, Australia

**Keywords:** Brain state, LSD, functional MRI, whole-brain modelling, perturbation, resting state networks

## Abstract

Lysergic acid diethylamide (LSD) is a potent psychedelic drug, which has seen a revival in clinical and pharmacological research within recent years. Human neuroimaging studies have shown fundamental changes in brain-wide functional connectivity and an expansion of dynamical brain states, thus raising the question about a mechanistic explanation of the dynamics underlying these alterations. Here, we applied a novel perturbational approach based on a whole-brain computational model, which opens up the possibility to externally perturb different brain regions in silico and investigate differences in dynamical stability of different brain states, i.e. the dynamical response of a certain brain region to an external perturbation. After adjusting the whole-brain model parameters to reflect the dynamics of functional magnetic resonance imaging (fMRI) BOLD signals recorded under the influence of LSD or placebo, perturbations of different brain areas were simulated by either promoting or disrupting synchronization in the regarding brain region. After perturbation offset, we quantified the recovery characteristics of the brain area to its basal dynamical state with the Perturbational Integration Latency Index (PILI) and used this measure to distinguish between the two brain states. We found significant changes in dynamical complexity with consistently higher PILI values after LSD intake on a global level, which indicates a shift of the brain’s global working point further away from a stable equilibrium as compared to normal conditions. On a local level, we found that the largest differences were measured within the limbic network, the visual network and the default mode network. Additionally, we found a higher variability of PILI values across different brain regions after LSD intake, indicating higher response diversity under LSD after an external perturbation. Our results provide important new insights into the brain-wide dynamical changes underlying the psychedelic state - here provoked by LSD intake - and underline possible future clinical applications of psychedelic drugs in particular psychiatric disorders.

**Highlights:**

- Novel offline perturbational method applied on functional magnetic resonance imaging (fMRI) data under the effect of lysergic acid diethylamide (LSD)
- Shift of brain’s global working point to more complex dynamics after LSD intake
- Consistently longer recovery time after model perturbation under LSD influence
- Strongest effects in resting state networks relevant for psychedelic experience
- Higher response diversity across brain regions under LSD influence after an external in silico perturbation

## 1. Introduction

In the past few years, we have witnessed an increasing interest in the study of the effects of psychedelic drugs, including lysergic acid diethylamide (LSD), on the human brain. LSD is a potent psychoactive drug, which was first synthesized in 1938 and whose potent psychological effects were discovered in 1943^1^. Between the 1950s and the late 1960s LSD was widely used in psychology and psychotherapy and its clinical applications as a pharmacological substance were well studied^2,3^, for a recent review and meta-analysis see Fuentes et al^4^. Due to political reasons and its widespread uncontrolled recreational use, LSD was made illegal in the late 1960s, which explains the hiatus period in human research with LSD. It was not until recently that the drug has undergone a renaissance in clinical and brain research.

Within the last few years, a significant number of human neuroimaging studies have been performed by only few research groups to identify neural correlates of the psychedelic state provoked by hallucinogenic drugs^5–11^. A non-exhaustive summary of these findings include: an increase in visual cortex blood flow and an expanded visual cortex functional connectivity^6^, a reduction of the integrity of functional brain networks^6,8,11^, a global increase in connectivity between networks^6,8^, where especially high-level association cortices comprising parts of the default-mode, salience, and frontoparietal attention networks and the thalamus showed increased global connectivity^8^, and an expanded repertoire of dynamical brain states, characterized by an increase of the variance of the Blood-Oxygen Level Dependent (BOLD) signal measured with functional Magnetic Resonance Imaging (fMRI) and a higher diversity of dynamic functional connectivity states^7^. While these results offer valuable insights into the major functional alterations taking effect in the brain during the psychedelic state, we do not yet have a compelling and complete mechanistic understanding of these effects in the context of whole-brain dynamics. To address this knowledge gap, we here apply a novel method combining a whole-brain computational model with an in silico model perturbation, previously described by Deco et al.^12^, which enables the simulation of external perturbations of any brain region for an unlimited amount of time in ways experimentally not possible.

In the last 15 years, there have been a number of studies investigating brain function by systematically exploring the dynamical responses to controlled artificial external perturbations of different brain regions and combining them with whole-brain neuroimaging^13–18^. There is a wide range of perturbation possibilities available, from easier to perform perturbation methods such as sensory stimulation and task instructions, to more invasive and costly methods, such as transcranial magnetic stimulation (TMS) in healthy human subjects to deep brain stimulation (DBS) in patients^19–22^. Also pharmacological studies inducing an anaesthetic state, which can also be considered as a perturbation to the brain, exist in human^23^ as well as in the non-human primate^24^ exploring the dynamic repertoires of the brain. The advantage of direct controlled artificial perturbations of specific brain regions is the systematic exploration of the provoked dynamical responses. These direct approaches have, however, been limited to transcranial magnetic stimulation (TMS) in healthy human subjects and to deep brain stimulation (DBS) in patients^19–22^.

Here we apply a novel in silico model perturbation approach to study the perturbation-elicited changes in global and local brain activity and to obtain a deeper understanding of the mechanisms underlying the experimentally observed dynamical brain changes under the influence of LSD in three different scanning conditions (rest, rest while listening to music and rest after listening to music). Previous studies have shown that the effects of LSD are amplified during listening of music^9,25,26^. Music is believed to act in combination with psychedelic drugs to enhance its emotional effects^25^ and that it acts synergistically with the drug to intensify mental imagery and access to personal memories^25,27,28^. We used a computational whole-brain model, which directly simulates the resting state BOLD signal fluctuations^12,18,29–31^ by simulating the dynamics in each brain area with the normal form of a supercritical Hopf bifurcation. This direct simulation of the resting state BOLD signal allows for systematical perturbation of each brain region in silico without needing to perturb the brain activity explicitly, e.g. via TMS. This whole-brain model based perturbation approach has proven useful to reveal the changes in brain dynamics underlying sleep, where brain activity was found to more rapidly return to its original state after perturbation than during awake^12^. Taken together with previous experimental findings on LSD, we hypothesized that under the influence of LSD, the brain would take longer to return to baseline activity - meaning brain activity without the model based perturbation - after a strong simulated perturbation. Such a scenario would be consistent with more complex and less stable dynamics^12,32^ as well as brain dynamics closer to bifurcation or critical regime^6–8,33^. Indeed, close to a bifurcation or instability, a dynamical system slows down its fluctuations and increases its responsiveness and complexity^31,34^. Whole-brain models have been shown to best represent the functional connectivity of whole-brain resting-state fMRI close to a bifurcation^31,34^. Previous research has suggested that LSD re-organizes brain dynamics at the edge of criticality^33^. Furthermore it has previously been shown that in an awake resting state - when compared to deep sleep - the brain takes longer to go back to its original state after perturbation^12^, and that perturbation induced stimuli propagate to other brain regions beyond the original stimulation site in an awake resting state as opposed to deep sleep^13,15,35^. Moreover, it has been shown that, while anesthesia reduces the complexity of brain signals with respect to normal wakefulness, LSD increases the activity complexity with respect to normal wakefulness, without a global loss of consciousness or changes in physiological arousal as seen in sleep or anaesthesia^36^. We thus hypothesized that LSD would produce more complex and sustained responses to perturbations than in normal resting-state conditions. We further expected this effect to be even stronger in the music condition, where the effects of LSD have been found to be amplified^9,25,26^.

## 2. Materials and Methods

### 2.1. Functional magnetic resonance imaging (fMRI) data

For the fMRI blood oxygen level dependent (BOLD) data, 20 healthy participants were scanned in 6 different conditions: LSD resting state, placebo (PCB) resting state, LSD and PCB resting state while listening to music, LSD and PCB resting state after listening to music. LSD and PCB sessions were separated by at least 14 days with the condition order being balanced across participants, who were blind to this order. All participants gave informed consent. The experimental protocol was approved by the UK National Health Service research ethics committee, West-London. Experiments conformed with the revised declaration of Helsinki (2000), the International Committee on Harmonization Good Clinical Practice guidelines and the National Health Service Research Governance Framework. The data collection was sponsored by the Imperial College London, which was carried out under a Home Office license for research with schedule 1 drugs. Eight out of the 20 subjects were excluded from further analyses for the following reasons: the scanning session of one participant needed to be terminated early due to the subject reporting significant anxiety. Four participants were excluded due to high levels of head movement (as described in the original publication by Carhart-Harris^6^, the exclusion criterion for excessive head movement was subjects displaying more than 15% scrubbed volumes with a scrubbing threshold of FD = 0.5). Three participants needed to be excluded due to technical problems with the sound delivery in the music condition. In total, 12 subjects were considered for further analyses. Each participant received either 75 g of LSD (intravenous, I.V.) or saline/placebo (I.V.) 70 minutes prior to MRI scanning. As described in the supplementary information of the original publication by Carhart-Harris et al^6^ the participants reported noticing subjective drug effects between 5 to 15 minutes post-dosing. The drug effects reached peak intensity between 60 to 90 minutes post-dosing. The subsequent plateau of drug effects varied among individuals regarding their duration, but participants reported a general remaining of the drug effects for four hours post-dosing. MRI scanning started - as mentioned above - approximately 70 minutes post-dosing, and lasted for about 60 minutes. After each of the three scans, participants performed subjective ratings inside the scanner via a response box. The subjects who received saline/placebo were considered as baseline MRI scans compared to the LSD scans. The BOLD fMRI data were recorded using a gradient echo planer imaging sequence, TR/TE = 2000/35ms, field of view = 220mm, 64×64 acquisition matrix, parallel acceleration factor = 2, 90° flip angle. The exact length of each of the BOLD scans per participant was 7:20 minutes. As described in the original publication by Carhart-Harris^6^, the performed pre-processing steps were the following: 1) the first three volumes were removed; 2) de-spiking; 3) slice time correction; 4) motion correction by registering each volume to the volume most similar to all others regarding least squares; 5) brain extraction; 6) rigid body registration to anatomical scans; 7) non-linear registration to 2mm MNI brain; 8) scrubbing using an FD threshold of 0.4 (the mean percentage of volumes scrubbed for placebo and LSD was 0.4 ±0.8% and 1.7 ±2.3%, respectively). The maximum number of scrubbed volumes per scan was 7.1% and scrubbed volumes were replaced with the mean of the surrounding volumes. Additional pre-processing steps were: 9) spatial smoothing of 6mm; 10) band-pass filtering between 0.01 to 0.08 Hz; 11) linear and quadratic de-trending; 12) regressing out 9 nuisance regressors (all nuisance regressors were bandpass filtered with the same filter as in step 10.

BOLD signals were averaged over cortical and sub-cortical regions of interest following the automated anatomical labeling (AAL) atlas parcellation of the brain into 90 regions of interest (76 cortical and 14 subcortical regions, AAL90), comprising 45 regions in each hemisphere^37^. We chose this parcellation of the human brain, since especially for studying the spatiotemporal dynamics on a whole brain level, AAL seems to be particularly well fitted. It has been found to produce good results in the whole-brain literature in general^12,38–40^ and furthermore whole brain computational models can be quite computationally expensive to perform and thus profit from a not too large number of parcels, as is the case in the AAL parcellation. The list of AAL ROIs can be found in the Supplementary Material (Supplementary Table S1).

The full details on the whole study design, the scanning protocol and further details on the fMRI pre-processing can be consulted in the supplementary information of the original publication^6^.

### 2.2. Anatomical connectivity

The anatomical connections between the different brain areas used in this study were obtained from Diffusion Tensor Imaging (DTI) data of an independent set of subjects, recorded in 16 healthy right-handed participants (11 men and 5 women, mean age: 24.75 ± 2.54), recruited through the online recruitment system at Aarhus University. This data has already been described in previous studies^29,41^. Briefly, the automated anatomical labelling (AAL) template was used for the parcellation of the entire brain into 90 regions, as explained in the previous section. The brain parcellations were conducted in each individual’s native space. The acquired DTI data was used to generate the structural connectivity (SC) maps for each participant. A three-step process was applied to construct these structural connectivity maps. First, the regions of the whole-brain network were defined with the AAL template as used in the functional MRI data. Secondly, probabilistic tractography was applied to estimate the connections between nodes in the whole-brain network (i.e. edges). Finally, the data was averaged across participants.

### 2.3. Hopf computational whole-brain model

The brain activity in each brain region was simulated with a computational whole-brain model, which has been previously described in various publications^12,29,31,42^. The model is based on the 90 coupled brain regions, comprising cortical and subcortical areas, retrieved from the AAL parcellation explained above. This computational model simulates the spontaneous brain activity in each node, which originates in the mutual interactions between anatomically connected brain areas (Fig. 1A). The anatomical connections are represented by the structural connectivity matrix *C_ij_*, obtained through DTI based tractography, as explained above. The structural connectivity matrix was scaled to a maximum value of 0.2^29,31^, leading to a reduction of the parameter space to search for the optimal parameter. The dynamics in each brain area can be simulated by the normal form of a supercritical Hopf bifurcation, which can describe the transition from noise-induced oscillations to fully sustained oscillations^31,43^. In fact, it has been shown that by coupling the brain regions together using the underlying anatomical connections, the interactions between the local Hopf oscillators can describe electroencephalography (EEG)^44^, magnetoencephalography (MEG)^41^ and fMRI dynamics^12,29,31,42^. The dynamics of a given uncoupled node *j* are described by the following complex-valued equation, representing the normal form of a supercritical Hopf bifurcation:

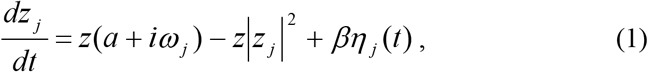

where 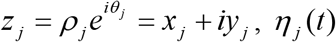, is additive Gaussian noise, *β* = 0.04 and *ω_j_* is the intrinsic node frequency, which was estimated as the peak frequency of the filtered BOLD time series for each brain region averaged over the participants within one subject group for each of the 6 conditions. This normal form possesses a supercritical Hopf bifurcation at *a* = 0. For *a* > 0 the local dynamics settle into a stable limit cycle, producing self-sustained oscillations with frequency 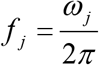. For *a* < 0 the damped oscillations lead the system to a stable fixed point (or focus), at *z_j_* = 0, and, in the presence of noise, noise-induced oscillations are observed.

**Figure 1:**
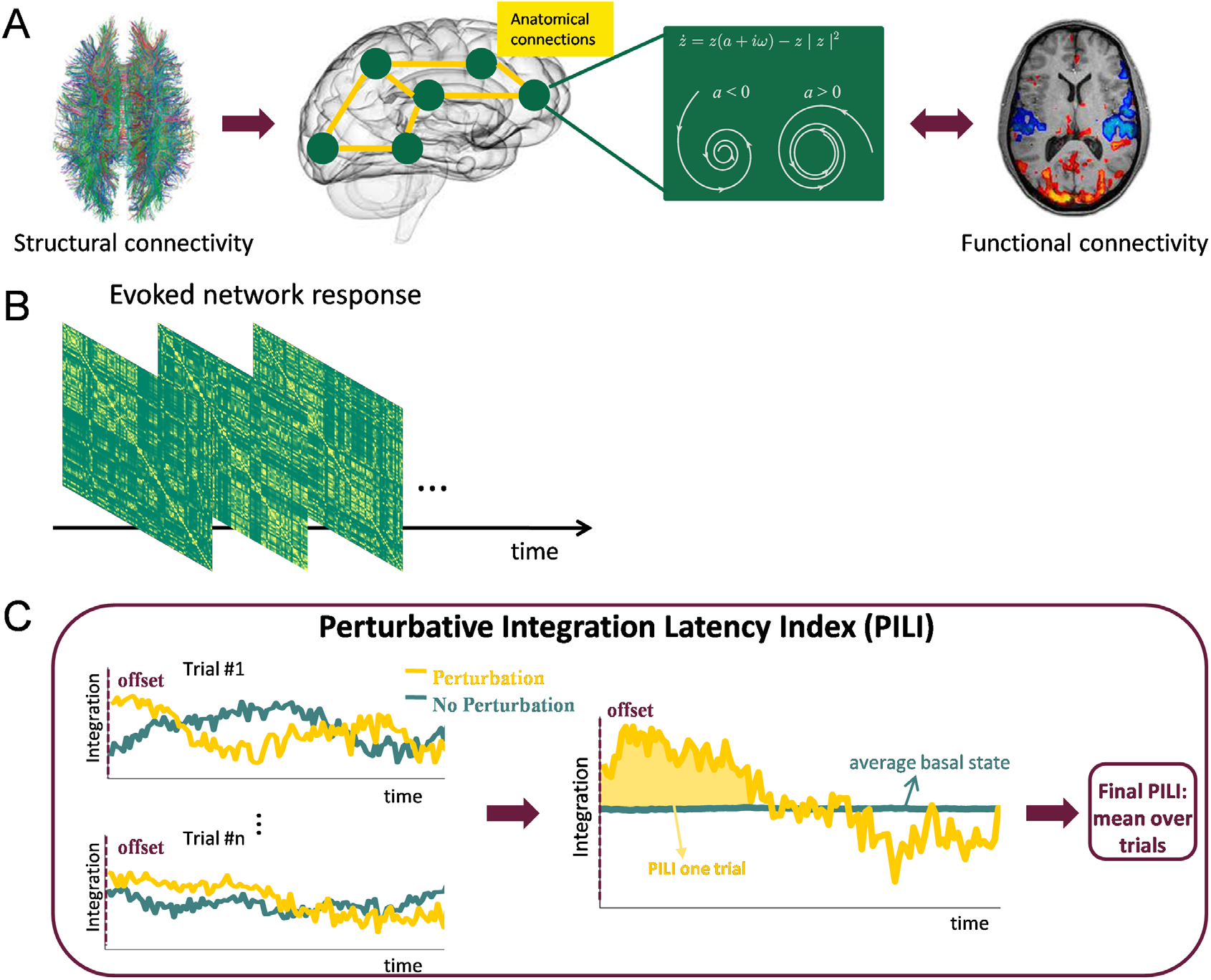
Calculation of the Perturbative Integration Latency Index (PILI). **A.** Initially, the computational whole-brain model was built based on the empirical structural connections between the 90 brain nodes. In this model each brain area was represented by a supercritical Hopf bifurcation. The model was fitted to the empirical functional connectivity in each of the 6 conditions, thus resulting in an optimal global coupling parameter for each condition. **B.** Next, we simulated the BOLD time series in each brain node for the basal dynamics and for the two perturbed states. The signals were band-pass filtered and Hilbert transformed to obtain the instantaneous phases and to subsequently calculate the phase locking matrix for each time point. **C.** Next, the integration was calculated as a function of time over 200 seconds in the basal state and after the offset of a model perturbation in either the synchronous or the noisy regime (here only shown the synchronous regime). The integration was computed by binarizing the phase locking matrix for different thresholds and calculating the number of areas in the largest connected component and finally integrating over thresholds. Finally the PILI was calculated, which characterizes the return of the brain dynamics to the basal state after a model perturbation of the system. For each trial, the PILI was computed as the integral under the curve of integration values after the offset of the model perturbation (yellow) until reaching the maximum of the basal state (blue). The final PILI was obtained by averaging over trials. (see section *2*. *Methods* for detailed explanation).

In order to simulate the whole-brain dynamics a coupling term was added which represents the input from node *j* to node *i* scaled by the structural connectivity matrix *C_ij_*. Hence, the whole-brain dynamics are described by the following set of coupled equations:

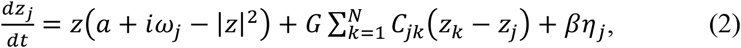

This model can be interpreted as an extension of the Kuramoto model^30,45^ with amplitude variations, hence the choice of coupling (*z*_*k*_ – *z*_*j*_), which relates to a tendency of synchronization between two coupled nodes. For each node *j* the variable *x_j_* = Re(*z_j_*) simulates the fMRI BOLD signal using the Euler algorithm with a time step of 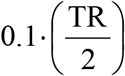. The parameter *G,* the global coupling strength, serves as a global coupling factor scaling equally the total input in each brain node.

### 2.4. Functional connectivity estimation

The BOLD signal of each AAL region was detrended, demeaned and then band-pass filtered within the range of 0.04-0.07 Hz following Glerean et al.^46^ individually for each subject. This frequency band has been shown to be less affected by noise and to be more functionally relevant compared with other frequency bands^46–49^. Next, the filtered time series were z-scored for each subject. The functional connectivity (FC) matrices were then calculated for each participant in each condition. Here we calculated the FC matrix as the Pearson correlations between the BOLD signals of all pairs of regions of interest (ROIs) over the whole recording duration. To obtain group-level FC matrices we applied fixed-effect analysis by Fisher’s r-to-z transforming (*z* = tanh(*r*)) the correlation values before averaging over all participants within each condition and then back-transforming to correlation values. Thus, we obtained 6 final FC matrices, one for each condition. For the group level comparison, the FC matrices were averaged across subjects individually for each condition and the comparison was performed for each pair of LSD - PCB scanning condition (i.e. LSD vs. PCB in rest, rest with music and rest after music conditions, respectively). To test the significance of the differences of the conditions, we generated 100 surrogate datasets where the LSD and PCB conditions are randomly permuted with a 50% chance of switching of the condition assignment, following Jobst et al.^29^. In this way, the group pairs get randomly mixed and thus fulfil the null-hypothesis of no difference between drug-induced conditions.

In order to ensure that within the group PCB there would be no differences between FC matrices between the group of participants who received PCB in their first session and the group of participants who received PCB in their second session, we performed a similar statistical significance analysis as described above. We divided the PCB sessions in the aforementioned groups and generated again 100 surrogate datasets where the group assignments are randomly permuted with a 50% chance of switching the group assignment. Thus, also here the null-hypothesis of no difference between the two groups is fulfilled and it can be analyzed if the differences of the mean FC matrices of the two groups are significantly larger than the ones generated by the surrogate data. The results of this analysis are shown in the Supplementary Material (Supplementary Figure S1).

In line with this analysis we furthermore analyzed if the differences between the LSD and PCB states showed differences between the two groups mentioned above, those who received PCB in their first session (“First”) and those who received PCB in their second session (“Second”). We again divided the data into these two groups and now compared the LSD state to the PCB state within each group, as was done in the original FC matrix analysis described above. Then, we analyzed the differences between the two groups “First” and “Second” regarding the differences between LSD and PCB states, a difference of differences so to speak. In order to test for statistical significance we again constructed surrogate data in the same fashion as described above and tested for significance. The results of this analysis can be consulted in the Supplementary Material (Supplementary Figure S2).

### 2.5. Drug state classification with Gaussian classifier

To establish how specific each of the functional connectivity matrices is to the drug state (LSD or PCB), we classified the drug state based on the covariance of fMRI signals using a jackknife cross-validation approach, assuming that observations are drawn from a multivariate Gaussian distribution, following Jobst et al^29^. First, we estimated the covariance (∑_LSD_ and ∑_PCB_) using the data of *n* −1 participants (train set), where *n* is the number of participants, for each drug state. Note, that in the Gaussian approximation the fMRI signals were fully determined by their covariance, since the data was z-scored and thus the mean of each fMRI time-series was zero. Then, we associated the data of the remaining subject (test set) to a drug state by selecting the zero-mean multivariate Gaussian process (*N* (0, ∑_LSD_) or *N* (0, ∑_PCB_)) which maximises the log-likelihood of the test data given the trained model. We calculated the percentage of correct classifications across both states and the *n* participants. Given the zero-mean multivariate Gaussian process *N* (0, ∑), the likelihood of a test N-dimensional vector *X_t_*, representing the *t*-th time step of the test data, is given by:

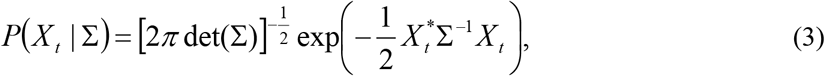

where det(∑) is the determinant of the covariance ∑ and the superscript * represents the transpose. The log-likelihood *L* of the entire test time series *X* = *X*_1,…,*T*_, where *T* is the number of time steps, is given by (assuming independence of the observations):

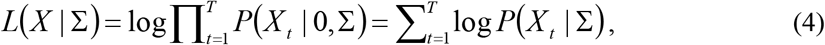

To summarize, we calculated *L*(*X* | ∑_LSD_) and *L*(*X* | ∑_PCB_) for each test dataset *X*. We predicted the state LSD if *L*(*X* | ∑_LSD_) > *L*(*X* | ∑_PCB_), otherwise the predicted state was PCB.

To assess statistical significance of the classification performance we computed the probability of obtaining *k* correct classifications by chance: 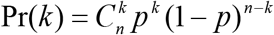, where *p* is the probability of getting a correct classification by chance 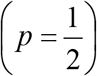 and *n* is the number of tests. Significant decoding of the conditions was reached when the performance exceeded the 95th percentile of Pr(*k*).

### 2.6. Fitting the model to experimental data

We explored the parameter space of the whole-brain computational model by varying the global coupling strength parameter *G* from 0 to 2 in steps of 0.01.To match the procedure applied on the empirical data, we filtered the simulated BOLD time series as well in the range of 0.04-0.07 Hz. Furthermore, the signal lengths of the simulated data coincided with the duration of the empirical data recordings. Next, the FC matrix was estimated on the simulated data for the whole parameter space applying the same procedure as on the empirical data. Then, the fitting between the empirical and the simulated FC matrices was calculated for each condition (i.e. LSD rest after music and PCB during rest, rest with music and rest after music) for the whole parameter space using the Kolmogorov-Smirnov distance (KS distance) between the two matrices, yielding a measure of fit for each value of the parameter *G* for each condition. For each condition, 50 simulations of the BOLD time series were generated, and the KS-distance of fit was averaged across the 50 simulations in order to minimize the random effects induced by the Gaussian noise in the model. We compared the resulting fitting curve minima with the surrogate data explained above in order to test for significant differences between the LSD and PCB conditions. The coupling parameter values, where the fitting curves were minimal, were then used for the following analysis steps.

### 2.7. Model perturbation protocols

Following Deco et al.^12^ we made use of the locally defined bifurcation parameter *a* of the Hopf model to simulate two kinds of off-line perturbation protocols evoking either deviations from the basal state (*a* = 0) into the synchronous regime (*a* > 0) or into the noisy regime (*a* < 0). In order to investigate the local effects provoked by the perturbation of single brain areas, we perturbed each node individually, repeated the perturbation procedure 3000 times and performed statistical analyses using the error of the distribution averaged over the 3000 trials. One perturbation trial consisted in perturbing one out of 90 nodes for 100 seconds by setting its local bifurcation parameter value *a* to either *a* > 0 or *a* < 0. Specifically, for the synchronization perturbation protocol *a* was set to 0.6 and for the noise perturbation protocol to −0.6. This leads to more oscillations in the perturbed node in the synchronization case and to an artificial destruction of the basal synchronization between the perturbed node and the other brain areas in the noise case. After perturbation, the bifurcation parameter was reset to zero in the perturbed node.

### 2.8. Integration measure

Next, in order to measure the level of brain-wide simulated BOLD signal interactions over time, we applied a measure previously defined in Deco et al.^50^ and applied to fMRI data in Deco et al.^12^, which characterizes the level of integration across all brain regions for each time point.

First, the Hilbert transform was applied on the band-pass filtered simulated time series giving us the instantaneous signal phases *φ_n_*(*t*). Next, the phase locking matrix *P* was calculated which characterizes for each time point the pair-wise phase synchronization between two brain regions *p* and *q*:

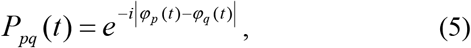

where *i* is the imaginary unit (Fig. 1B). The level of integration at time *t* is then defined as the size of the largest connected component of the phase locking matrix averaged over thresholds^12,50^. We binarized the phase locking matrix *P* for 100 evenly spaced thresholds between 0 and 1, applying the criterion |*P*| < *θ* = 0 and 1 otherwise, and extracted for each of the thresholds the number of nodes of the largest connected component of *P*(*t*) at each time point *t*. We then calculated the integration *I* (*t*) at time *t* as the integral of the curve of the largest component as a function of the thresholds (Fig. 1C). We computed the integration over 200 seconds of simulated BOLD time series in the basal state and starting at perturbation offset in the perturbed case.

### 2.9. Perturbative Integration Latency Index (PILI)

Following Deco et al.^12^ we calculated the Perturbative Integration Latency Index (PILI) to characterize the return of the brain dynamics to the basal state after a model perturbation of the system (Fig. 1C). For this we used the changes of the level of integration over time from the perturbed state to the basal dynamics.

First, the integration was calculated for 200 seconds of the simulated basal state (blue curve in Fig. 1C), averaged over 3000 trials and finally the maximum and minimum values of the averaged curve were identified. This was done for each of the 6 conditions. Then, the system was perturbed following the procedure described above and again the integration was computed over 200 seconds after the offset of the perturbation. This procedure was performed 3000 times. The maximum and minimum values of the basal integration curve were used to determine the moment of recovery after the model perturbation, for the synchronization and noise protocol, respectively. Then, the PILI was calculated as the integral of the integration curve from perturbation offset to the reaching point of the basal state. Finally, we computed the average PILI over trials to obtain one final value for each brain area. The PILI reflects how strong the system reacts to a model perturbation and how long it takes for it to regain its basal dynamical state. The statistical significance tests were performed across the 3000 trials applying a Mann-Whitney U test to compare between LSD and PCB induced states.

### 2.10. Region-wise and resting state network analysis

The above described procedure resulted in one PILI for each of the 90 brain areas. We compared the p-values for all brain regions between LSD and PCB in each of the three scanning conditions (rest, rest with music, rest after music), computed with the above described statistical significance test, after ordering them from smallest to largest. Bonferroni correction was applied in order to correct for the multiple comparisons across the 90 brain areas.

Next, we evaluated the differences between PILI values in seven commonly observed resting state networks (RSNs): default mode network (DMN), executive control, dorsal attention, ventral attention, visual, limbic and somato-motor networks, as described in Yeo et al.^51^. The parcellation of the cerebral cortex into these 7 networks has been extracted from the intrinsic functional connectivity data from a group of 1000 participants^51^ and is available online at http://surfer.nmr.mgh.harvard.edu/fswiki/CorticalParcellation_Yeo2011. For each of the 7 RSN^51,52^, we computed the standardized difference between the PILI values in the LSD and PCB induced states by calculating Cohen’s d-values, defined as 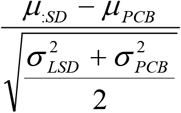, where μ is the mean of the PILI values and σ the standard deviation,^53^ by taking into account only the brain areas belonging to that particular RSN. The RSNs were then ordered from highest to lowest Cohen’s d-value, where the higher the d-value, the higher the difference between PILI values and thus the larger the response to a model perturbation under the influence of LSD in one particular RSN. For completeness, we furthermore tested for statistical significance between the LSD and the PCB state models in each condition for each RSN by applying a Mann Whitney U test on the final PILI values of the brain areas belonging to each particular RSN. Bonferroni correction was applied to correct for the multiple comparisons across the 7 RSNs.

### 2.11. Response variability

Finally, in order to learn more about the differences between the dynamics of individual brain regions, we calculated the variability of the PILI values over different brain regions. This was done by calculating the standard deviation of the PILI values across all brain nodes for each of the 3000 trials and then comparing the distributions over trials between LSD and PCB brain state model. We evaluated statistically significant differences between the LSD and PCB induced brain states by applying a two-sided t-test.

## 3. Results

We investigated the differences between LSD and PCB brain states in three different scanning conditions, namely LSD and PCB during rest, LSD and PCB during rest while listening to music and LSD and PCB during rest after listening to music. We applied a previously published off-line perturbational approach based on a whole-brain model, which characterizes the return of the brain dynamics to the basal state after a model perturbation of the system (see Fig. 1 for overview of the method).

### 3.1. Functional connectivity and optimal working point

Firstly, we investigated the differences in functional connectivity (FC) between the LSD and PCB brain states in all three scanning conditions. For this, we calculated the FC matrices on a subject-level basis and averaged across subjects within each condition (see section *2*. *Methods*). To compute the differences between the LSD and PCB states, the mean FC value was computed for each condition and then compared with the surrogate data. We found a significant difference in the mean FC values between LSD and PCB in the music condition (LSD: 0.204±0.179, PCB: 0.140±0.197; p-value: 0.0297). We also observed a slight increase in mean FC values during the LSD state with respect to PCB in resting conditions, which did not involve listening to music(rest: LSD: 0.186±0.175, PCB: 0.154±0.202; p-value: 0.0990; rest after music: LSD: 0.181±0.171, PCB: 0.163±0.191, p-value: 0.1485). However, these differences were not found to be significant (Fig. 2A).

**Figure 2:**
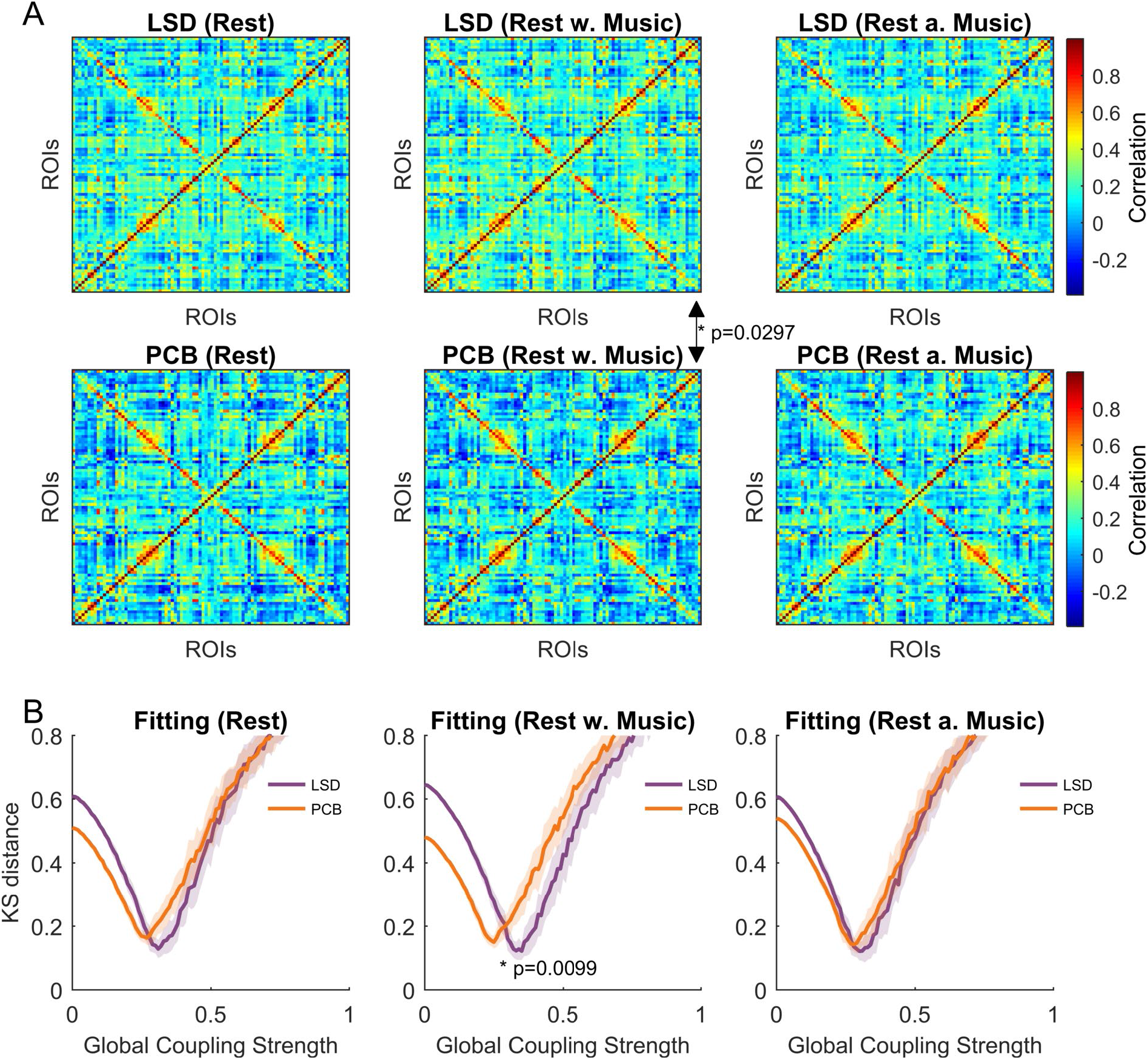
Empirical functional connectivity and model fitting. In **A** the functional connectivity matrices are shown for each of the 6 conditions. Significance tests have been performed between the LSD and PCB conditions resulting in a significant difference in the mean functional connectivity between the LSD and the PCB state in the music scanning session. In **B** the mean and standard deviation over 50 realizations of the KS distance between the empirical and the simulated functional connectivity matrices are shown for each condition as a function of the global coupling strength. The optimal fit corresponds in each condition to the minimal KS distance. We found a significant difference between the optimal fit in the LSD and the PCB state in the music scanning session.

Next, we fitted the Hopf whole-brain model to the fMRI data in each condition in order to compare the effects of LSD and PCB with regards to their dynamical working point, i.e. the parameter region where the model best fits the data. The Hopf whole brain model has been previously shown to be able to simulate fMRI-BOLD network dynamics^12,29,31,42^ and is especially well suited for simulating external perturbations to distinct brain nodes, as demonstrated in Deco et al.^12^. We computed the KS distance between the empirical and the simulated functional connectivity matrices and found a shift in the optimal global coupling parameter *G*, i.e. the minimal KS distance, towards higher values under the influence of LSD in all three scanning conditions (rest: LSD: 0.31, PCB: 0.27; rest with music: LSD: 0.35, PCB: 0.25, rest after music: LSD: 0.29, PCB: 0.28) with a significant difference in the music condition (p = 0.0099) (Fig. 2B). As above, to assess statistical significance, the values were compared with surrogate data obtained by randomly permuting group assignments (see section *2*. *Methods*).

To summarize, we found a global increase in functional connectivity and a shift of the optimal global coupling strength to larger values under the effect of LSD, implying a higher global level of brain connectivity in this state.

### 3.2. Drug state classification with Gaussian classifier

We assessed how specific the functional connectivity is to the drug state (LSD or PCB). The jackknife cross-validation procedure we applied consisted of: first, calculating the covariances on a subset of the data using N-1 participants, and then classifying the data of the remaining subject given the previously computed covariances (see *Methods*). We found that the drug states were predicted with an accuracy exceeding the significance level for all 3 scanning conditions (75% for rest, 79,17% for rest with music and 70,83% for rest after music) (Supplementary Figure S3). Importantly, these classification performances were significantly higher than expected by chance given the number of subjects. To summarize, the whole-brain covariance of single participants reliably relates to the drug state and thus even a small number of participants can be seen as representative of the two states LSD and PCB.

### 3.3. Global differences in Integration

Next, we simulated two kinds of model perturbation protocols for each brain state in order to compare the different state models with regard to their responses to a strong in silico perturbation. We compared the brain states by making use of the global integration measure (see section *2.7*. *Integration measure*), which we used to evaluate the differences in integration.

With the adjustment of the whole-brain model to the fMRI data, we obtained a representative model of the basal brain state for LSD and PCB states in each condition. The two model perturbation protocols were then simulated by either shifting one brain node to a more synchronous state or to a noisier state for 100 s (see section *2.6*. *Model perturbation protocols*). This was done for each of the 90 nodes representing the brain regions in the AAL parcellation. Immediately after perturbation, we quantified the perturbation-caused changes in brain-wide signal interactions over time by computing the global integration measure.

In Fig. 3, the integration averaged over 3000 trials and all 90 brain nodes is displayed as a function of time. The integration is shown immediately after perturbation offset for LSD and PCB state models in each condition. We observed that the basal integration was higher for each scanning condition in LSD (dark green curve) compared with PCB (light green curve), where the difference between LSD and PCB was highest in the music condition. This implies that without perturbation, the level of BOLD signal connectedness was higher in the LSD state than in PCB. Notably, comparison of the basal integration among scanning conditions (i.e. before, during and after music listening) within both the LSD and PCB state models also revealed that the basal integration increased under the influence of LSD while listening to music, whereas in the PCB state model it decreased with music. This finding is in line with previous results that have demonstrated an enhancement of the LSD experience while listening to music^9,25,26^, whilst in the PCB state, music appeared here to have a contrastive effect. These results call for further exploration of the differential effects of music on brain dynamics in the psychedelic state. Regarding the perturbation protocols, we found that for all three scanning conditions, the deviations from the basal activity were both stronger and longer-lasting under the influence of LSD (violet curve)in comparison with PCB (orange curve) after being exposed to the same kind of perturbation. While this is valid for both synchronization protocols and noise protocols, the effects on the differences in integration in the LSD state model as compared to the PCB state model were much smaller for the noise protocol than for the synchronization protocol (detailed analysis in *Methods - Global and local differences in Perturbative Integration Latency Index* and Supplementary Information). We therefore decided to mainly focus on the synchronization protocol for the rest of the article. The results of the noise perturbation protocol can be consulted in the Supplementary Information.

**Figure 3:**
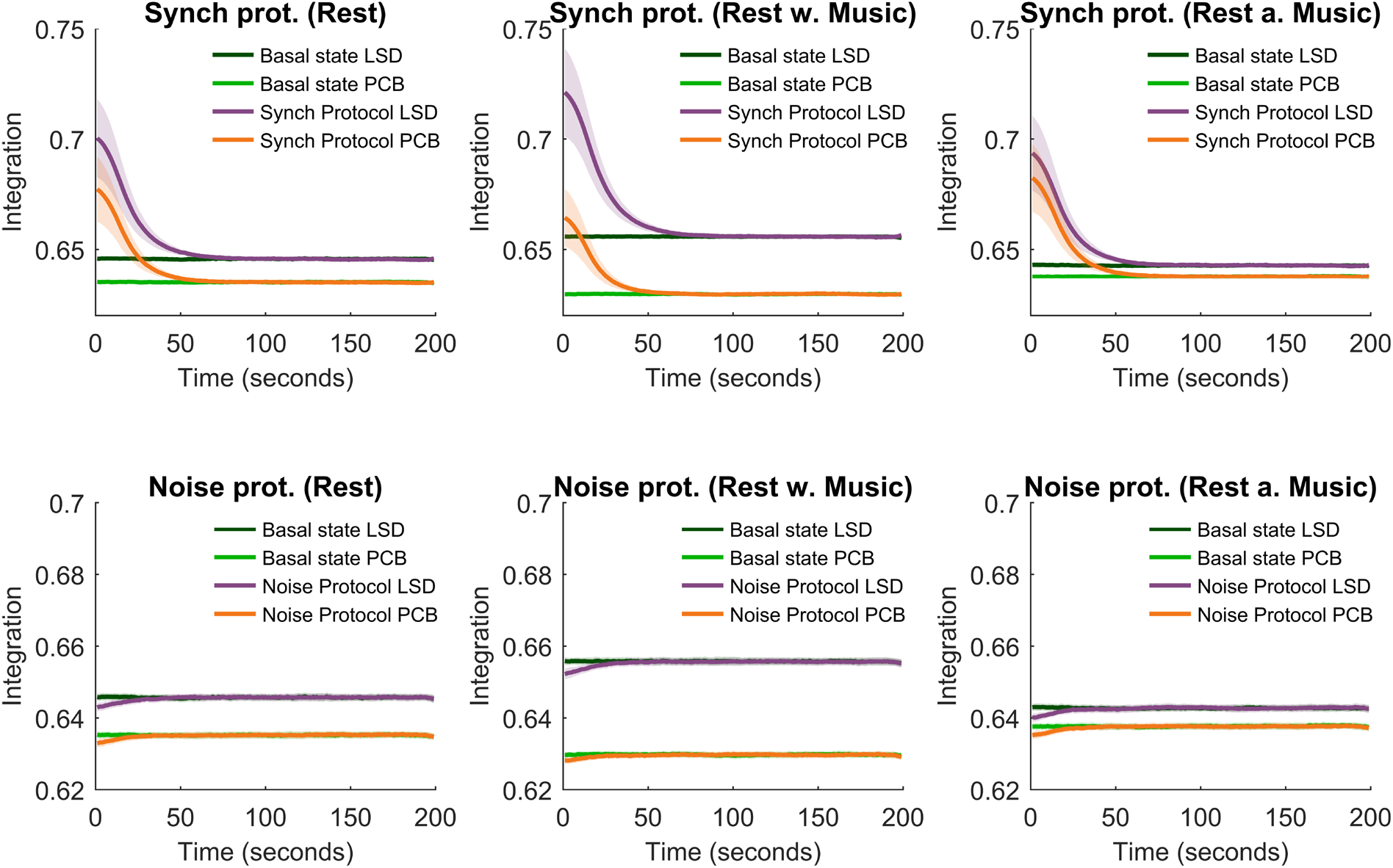
Mean integration. The integration averaged over trials and nodes and the standard deviation of the integration over nodes is shown as a function of time for the three scanning conditions for both perturbation protocols. The mean and standard deviation of the integration are shown in dark green and light green for the basal state of the LSD and the PCB state, respectively. The mean and standard deviation of the integration are indicated in violet and orange and for the LSD and the PCB state, respectively.

### 3.4. Global and local differences in Perturbative Integration Latency Index

In order to formally characterize the above observed changes in Integration strength and the return duration of the brain dynamics to its basal state after a model perturbation, we computed the Perturbative Integration Latency Index (PILI). The PILI is defined as the area under the integration curve up to the point it reaches the basal state. Thus, the PILI captures both, strength of deviation from the basal state and duration of the recovery. The PILI was calculated for each node by only perturbing this specific node and leaving the other nodes at their basal dynamics for 3000 trials, which were then averaged in order to obtain one single PILI value for each brain area (see section *2.8*. *Perturbative Integration Latency Index (PILI)*).

We found consistently higher PILI values for the LSD induced brain state model than for PCB in all three scanning conditions, where the effect was strongest for the music condition (Fig. 4). Again, the effect was diminished in the rest after music condition, which is most likely due to the decreased effect of LSD, as explained above. Most importantly, we demonstrate here, that the LSD and PCB brain states show very different dynamical responses to a model perturbation. In particular, the responses to the same perturbation are stronger and longer lasting under the influence of LSD with respect to PCB. Similar results were found for the noise protocol (Supplementary Figure S4). Also, here we observed a global increase in PILIs for LSD when compared to PCB for all three scanning conditions.

**Figure 4:**
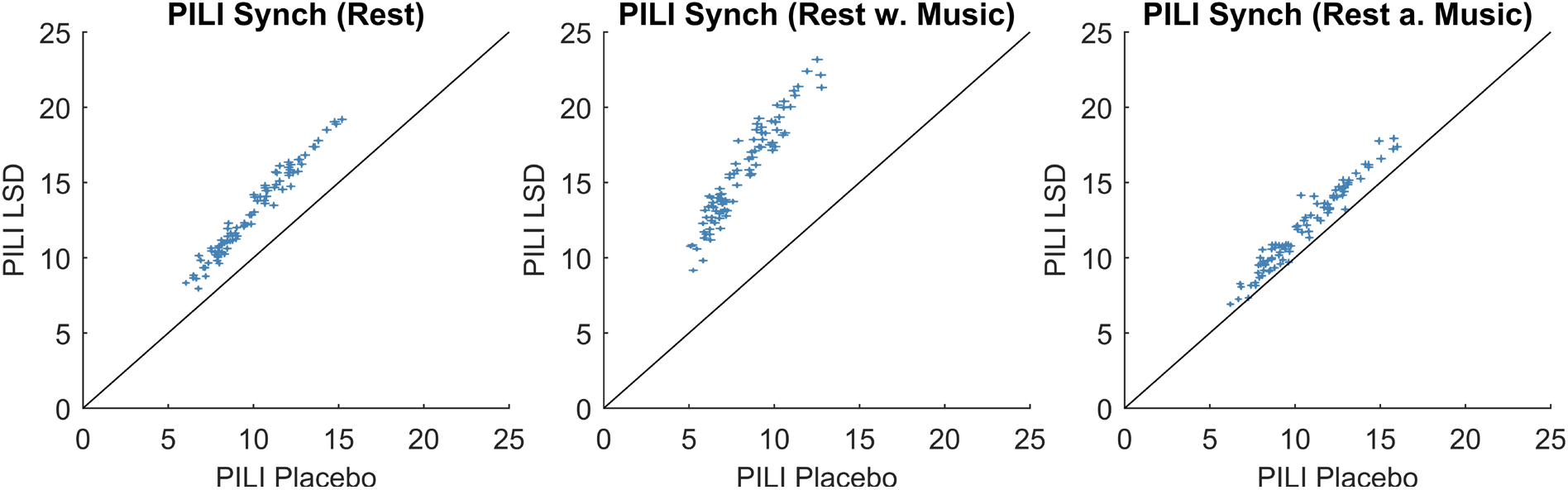
PILI - Node level analysis. Here the mean and the standard error of the mean (SEM) of the PILI values over trials are shown for each of the three scanning conditions for the LSD and the PCB state for all 90 brain regions. The vertical error bars represent the SEM for the PCB state and horizontal error bars represent the errors for the LSD state. The results show that the global differences between the LSD and PCB induced brain states were amplified in the music condition. Node-by-node analysis with corresponding p-values can be found in Table 1 and Supplementary Table S2.

**Table 1:** Node level PILI differences. In this table brain nodes are ordered for each scanning condition by p-values - from smallest to largest -, based on the PILI differences between LSD and PCB by perturbing each specific node at a time. Here the 20 regions with the smallest p-values are shown with their corresponding effect sizes.

| Rest |  |  | Rest with Music |  |  | Rest after Music |  |  |
| --- | --- | --- | --- | --- | --- | --- | --- | --- |
| Brain region | p-value | Effect Size | Brain region | p-value | Effect Size | Brain region | p-value | Effect Size |
| Olfactory R | 2.99e-37* | 0.2328 | Cingulum Mid R | 1.24e-172* | 0.5114 | Hippocampus R | 1.63e-34* | 0.2237 |
| Thalamus L | 2.70e-36* | 0.2297 | Precuneus L | 2.21e-166* | 0.5019 | Cingulum Ant R | 2.20e-21* | 0.1734 |
| Supp Motor Area R | 4.74e-35* | 0.2255 | Medial OFC R | 2.93e-166* | 0.5017 | Precuneus R | 4.97e-18* | 0.1580 |
| CingulumMid L | 3.34e-33* | 0.2192 | Frontal Sup Medial R | 1.32e-159* | 0.4915 | Precentral R | 4.12e-15* | 0.1433 |
| Calcarine L | 1.41e-32* | 0.2170 | Frontal Sup Medial L | 3.68e-158* | 0.4892 | Hippocampus L | 7.24e-12* | 0.1251 |
| Cingulum Ant R | 1.80e-31* | 0.2131 | Frontal Sup R | 9.98e-157* | 0.4870 | Supp Motor Area R | 9.00e-12* | 0.1245 |
| Occipital Sup R | 9.14e-30* | 0.2069 | Frontal Sup L | 1.24e-156* | 0.4868 | Occipital Mid L | 3.36e-11* | 0.1210 |
| Cingulum Post R | 1.03e-29* | 0.2067 | Precuneus R | 1.68e-154* | 0.4834 | Frontal Sup Medial R | 5.16e-11* | 0.1199 |
| Precuneus L | 2.19e-29* | 0.2055 | Cingulum Post L | 5.07e-151* | 0.4779 | Cingulum Mid L | 6.66e-11* | 0.1192 |
| Medial OFC L | 3.74e-29* | 0.2046 | Cingulum Mid L | 4.49e-149* | 0.4748 | ParaHippocampal R | 7.58e-11* | 0.1188 |
| Putamen L | 4.22e-29* | 0.2044 | Cingulum Post R | 6.79e-149* | 0.4745 | Medial OFC L | 2.80e-10* | 0.1152 |
| Thalamus R | 8.44e-29* | 0.2033 | Medial OFC L | 2.74e-147* | 0.4719 | Cingulum Ant L | 9.31e-10* | 0.1118 |
| Calcarine R | 2.85e-28* | 0.2013 | Caudate L | 2.64e-144* | 0.4670 | Frontal Sup R | 3.58e-09* | 0.1078 |
| Putamen R | 2.86e-28* | 0.2013 | Olfactory R | 1.63e-139* | 0.4591 | Fusiform R | 3.62e-09* | 0.1077 |
| Lingual L | 3.52e-28* | 0.2010 | Frontal Sup Orb L | 1.26e-136* | 0.4542 | Cingulum Mid R | 4.11e-09* | 0.1073 |
| Olfactory L | 1.10e-27* | 0.1991 | MedialOFC R | 1.22e-132* | 0.4474 | Calcarine R | 4.54e-09* | 0.1070 |
| Precuneus R | 1.12e-26* | 0.1952 | Cingulum Ant R | 5.57e-132* | 0.4463 | Temporal Pole Sup L | 1.10e-08* | 0.1043 |
| Cingulum Post L | 1.58e-26* | 0.1946 | Supp Motor Area R | 5.64e-131* | 0.4446 | Frontal Mid Orb L | 1.55e-08* | 0.1033 |
| Frontal Sup Medial L | 3.00e-26* | 0.1935 | Cingulum Ant L | 1.03e-130* | 0.4441 | Precuneus L | 2.14e-08* | 0.1022 |
| Cingulum Ant L | 3.87e-26* | 0.1931 | Frontal Sup Orb R | 5.13e-130* | 0.4429 | Temporal Inf L | 3.25e-08* | 0.1009 |

\* statistically significant after Bonferroni correction

In order to prove that the higher PILI values not only depend on the stronger deviations from the baseline brain activity, but are indeed longer lasting under the influence of LSD when compared to PCB, we furthermore calculated the time for the perturbed signals to come back to the basal state. We found, by applying a Mann Whitney U test, that for the synchronization protocol in the first resting state 88 out of 90 nodes showed significantly higher latencies in LSD when compared to PCB, in the Rest with Music condition 90 out of 90 nodes showed significantly higher latencies in the LSD state and in the Rest after Music condition 62 out of 90 nodes showed significantly higher latencies in the LSD state. This means that the perturbation effect is also longer lasting and not only stronger in the LSD state, where the effect is most prominent in the Music condition. The latencies for the 3 LSD conditions compared to the PCB conditions can be found in the Supplementary Material (Figure S5).

Next, to gain further insights into local processes, we looked at the PILI values on a node-to-node basis. We checked for statistical significance of the difference in the mean PILI value between LSD and PCB for each scanning condition for each node applying a Mann-Whitney U test with Bonferroni correction for multiple comparison across the number of brain nodes. The results for the synchronization protocol are shown in Table 1, where the 20 brain areas with the highest PILI differences are shown in order from smallest to largest p-value with their according effect sizes 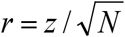, where *N* is the number of samples. Effect sizes between 0.1− < 0.3indicate small effects, 0.3− < 0.5 medium effects and ≥ 0.5 large effects. The ordering of the rest of the brain regions and the results for the noise protocol can be found in the Supplementary Material (Supplementary Tables S2 and S3).

Ordering the brain regions by p-values of each scanning condition revealed that globally p-values were lower and effect sizes higher for the rest with music condition with respect to the other resting conditions, which confirms previous findings on the amplified effect of LSD while listening to music^25,33^. The brain regions with small p-values in all three scanning conditions, were the cingulate cortex, the precuneus, the medial OFC and the supplementary motor area. Other regions where high differences between LSD and PCB could be observed were the calcarine sulcus, the olfactory sulcus, the superior frontal gyrus and the medial frontal gyrus, thalamus and hippocampus.

Taken together, these results reveal that the dynamical responses of the brain as a whole to an external model perturbation are stronger and longer lasting under the influence of LSD when compared to PCB. Furthermore, this effect is amplified in the model estimated from data in which participants listen to music. Next, we performed the same analysis on a resting state network level, in order to assess whether some networks exhibit larger responses to external perturbations than others and more importantly, whether those networks coincide with the ones which have been reported to be relevant for the LSD experience.

### 3.5. Relationship of PILI to resting state networks

Next, we assessed the differences in PILI values based on the synchronization protocol in seven reference RSNs - default mode, executive control, dorsal attention, ventral attention, visual, limbic and somato-motor networks - by computing Cohen’s d values, a standardized difference measure, between LSD and PCB PILI values for each RSN. Furthermore we tested for statistical significance of the differences between LSD and PCB state models for each RSN

The differences between LSD and PCB state models for all 7 RSNs in the resting state and music condition have been found to be statistically significant. In the rest after music condition 5 out of the 7 networks don’t survive the Bonferroni correction for multiple comparisons. The table of the corresponding p-values can be found in the Supplementary Material (Supplementary Table S4).

Notably, in all three scanning conditions, three RSNs were found to have the highest PILI differences between the LSD and PCB state models: i.e. the limbic, visual and default mode networks. The limbic network showed the highest differences in all three cases (see Fig. 5, where the RSNs were ordered for each of the three scanning conditions by Cohen’s d values, darker colours indicate higher difference). In both of the no-music conditions, the visual network seemed to play an important role, whereas in the music condition the default mode network showed higher differences in PILI values than the visual network. In the resting state conditions, the somato-motor network came fourth to the first three RSNs by Cohen’s d values, whilst in the music condition, the ventral attention network gained more importance.

**Figure 5:**
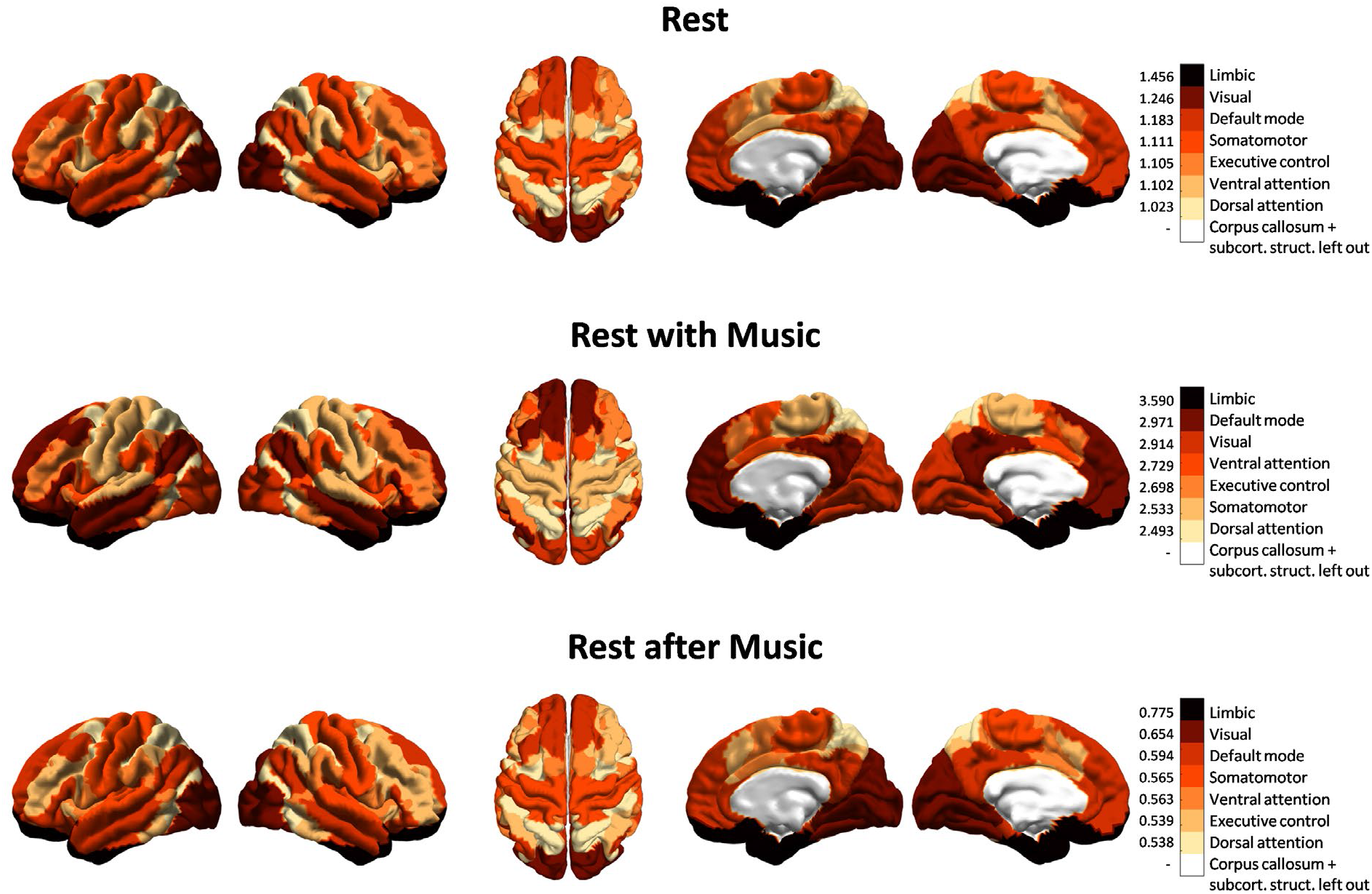
PILI - RSN analysis. The differences between the PILIs in LSD and PCB are shown on an RSN level. For all the nodes forming part of one RSN the Cohen’s d value was calculated based on the mean and standard deviation over nodes in each state, indicating the standardized mean difference between the PILIs of each RSN in LSD and PCB. This was done for each of the 7 RSNs. The RSNs were ordered for each scanning condition (rest, rest with music, rest after music) by Cohen’s d values, where darker colours indicate larger differences in PILI between the LSD and PCB conditions. The white area, which represents the corpus callosum and the subcortical structures, is to be discarded. It should be noted that the differences between PILI values in LSD and PCB state models for each RSN have found to be statistically significant in the rest and the rest with music condition. In the rest after music condition only 2 out of 7 networks (limbic network and DMN) show statistically significant differences (see Supplementary Table S4).

Overall, these results highlight that in particular three resting state networks, limbic, visual and default mode, show highly increased sensitivity under the influence of LSD, in line with previous studies^6,8^. Importantly, our findings propose a mechanistic explanation for the enhanced emotional, visual and self-referential processing due to increased sensitivity of the limbic, visual and default mode networks, respectively, in the psychedelic state.

### 3.6. Increased perturbation response variability in LSD condition

Finally, we analyzed the perturbation response variability across all brain regions. This was done by computing the standard deviation of the PILI values over brain nodes. In Fig. 6 we show the distribution over the 3000 trials of the standard deviation for all three scanning conditions and both drug states for the synchronization protocol. We found that the differences in variability between LSD and PCB were highly significant (p < 0.0001) in all three scanning conditions, with higher response variability under the influence of LSD than for PCB. This effect was strongest in the music condition and again less apparent in the after-music condition.

**Figure 6:**
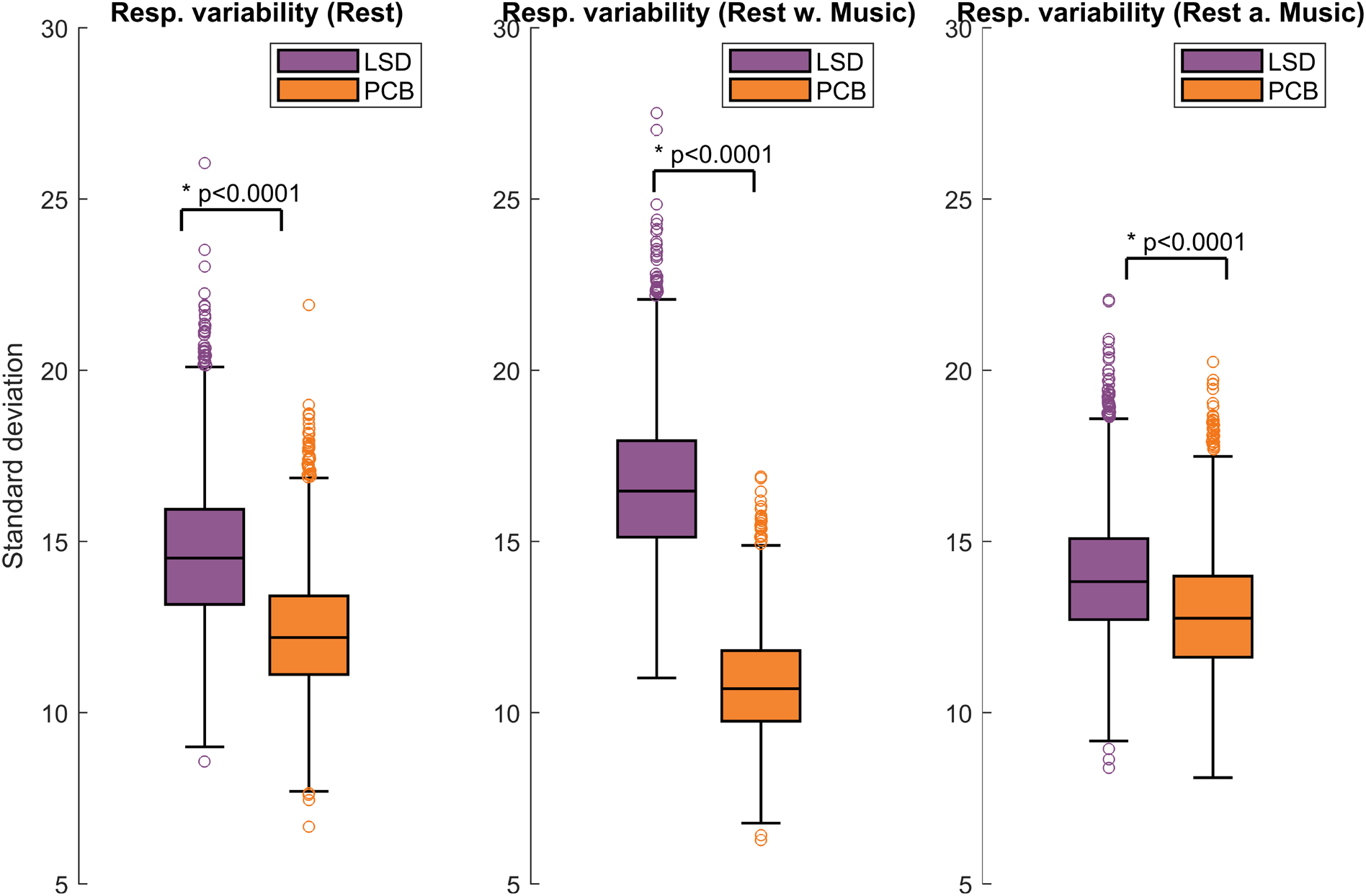
Response variability. Here the distribution over trials of the standard deviation of PILI values is shown for the three different scanning conditions for LSD and PCB. Statistical differences between LSD and PCB brain states were evaluated with a two-sided t-test resulting in highly significant differences in all three scanning conditions with significantly higher PILI variability in the LSD state with respect to PCB. Especially in the music condition under the influence of LSD a considerably larger response variability can be observed with a p-value significantly smaller than 0.0001.

## 4. Discussion

We applied a novel in silico model-based perturbational approach to analyze the perturbation-elicited changes in global and local brain activity under the influence of LSD compared with PCB in three consecutive scanning conditions, namely a resting state followed by resting while listening to music and finally a post-music resting state. Besides finding an increase in global functional connectivity and a shift of the brain’s global working point to higher connectivity in the LSD state, we showed that under the influence of LSD, brain dynamics show a larger divergence from and take longer to return to baseline activity after a strong model perturbation compared with the PCB state. Although we found that this effect was global on the whole cortex, our findings also revealed that certain brain regions and networks, such as the limbic network, the visual network, and the default mode network, were most sensitive to these changes. Finally, we also evaluated the differences between LSD and PCB with regard to the variability of these perturbational responses and found higher response variability under the influence of LSD.

We found that the empirical functional connectivity was higher on average in the LSD condition compared with the PCB condition, and this difference was especially pronounced in the music condition (Fig. 2A), where the effects of LSD seem to be amplified - as reported in the literature^9,25,26^. This finding consolidates the results of previous studies, where it was found that high-level association cortices and the thalamus exhibit increased global functional connectivity under the influence of psychedelics^8,54,55^. At least two previous studies have found increased thalamic functional connectivity to various cortical regions^54,55^ and another found a dramatic increase in functional connectivity between the primary visual cortex and other cortical areas under LSD -an effect that correlated strongly with ratings of enhanced visual imagery^6^. Similar results have been reported for other psychedelic drugs such as psilocybin (the main psychedelic component of magic mushrooms). One study found an expanded repertoire of dynamical brain states under the influence of psilocybin, characterized by an increase of the variance of the Blood-Oxygen Level Dependent (BOLD) signal measured with (fMRI) and a higher diversity of dynamic functional connectivity states^7^. In another study psilocybin was found to have an increasing effect on DMN-Task-positive network (TPN) functional connectivity, thus underlining similarities of the psychedelic state to psychosis and meditatory states, where the same effect has been found^54^. Yet another study by Roseman et al^56^ found an increase in between-network functional connectivity under psilocybin, suggesting that the psychedelic state makes networks become less differentiated from each other. All these findings confirm our results of an increase of global functional connectivity.

Additionally to comparing the functional connectivity between LSD and PCB, we also assessed how specific the functional connectivity is to the drug state, meaning how well the functional connectivity of a single participant relates to either the LSD or the PCB state. We found that the brain states were predicted with an accuracy exceeding the significance level for all 3 scanning conditions (see Supplementary Figure S3). The finding that the FC matrices of single participants can be classified to the corresponding drug state with an accuracy higher than the chance level, implies that the characteristics of the single subjects are reflected in the group-level results. Importantly, these classification performances were significantly higher than expected by chance given the number of subjects. This suggests that also a small number of participants, as is the case in this study, and the characteristics of their fMRI recordings for each of the two drug states can be seen as a representative sample which can be used to draw general conclusions on a global level. Nevertheless, it would be undoubtedly advantageous to perform further similar experiments in the future with more participants involved.

In order to study the whole-brain dynamics underlying the psychedelic state, first, we applied a whole-brain model based on the normal form of a supercritical Hopf bifurcation simulating directly the fMRI BOLD responses. Our analyses revealed that the global working region of brain dynamics shifts to higher global coupling parameters in the LSD state when compared with PCB. Notably however, statistical significance was only reached in the music condition, implying that the differences in brain dynamics between the LSD and PCB state may be accentuated under conditions of significant emotional evocation here represented by listening to music (Fig. 2B). This result underlines yet again the enhancing effect of music on the psychedelic state, as previously reported^9,25,26^. Taken together, our results suggest increased propagation of activity and enhanced communication between distinct brain regions. This finding is in agreement with previous studies that have demonstrated that the dynamical repertoire of the brain expands under the influence of psilocybin^7^, implying that, in this state, the brain operates in a different dynamic working region. Similar findings have also been recently demonstrated by Atasoy et al.^33^, where LSD was found to tune brain dynamics closer to criticality, entailing an increase in the diversity of the repertoire of brain states – a finding replicated more recently using both LSD and psilocybin data^57^. Increased brain criticality is consistent with the so-called entropic brain hypothesis^58,59^- and note the schematic figure 2 in Carhart-Harris et al.^32^.

In order to understand the optimal working point of brain dynamics in each scanning condition, we evaluated the responses to strong off-line model perturbations in each state. In a previous study^12^, this method was successfully used to discriminate between awake and sleep states. The importance of this new methodology lies in the fact that perturbations are exclusively applied in silico to a whole-brain computational model, allowing for stronger, longer lasting and brain node-specific perturbations in ways not possible experimentally. Furthermore, an important difference of this model-based perturbation approach to previously described perturbation procedures^13,15,35^ is the fact that with this new approach, we measure the recovery characteristics of the system after the offset of the perturbation, not the dynamical reaction to the perturbation itself.

Following this approach, we characterized return to the basal brain activity by the Perturbative Integration Latency Index (PILI). Interestingly, we found differences in the global integration, even without applying any perturbation, where the basal integration was increased under LSD in contrast to PCB, which was again amplified in the music condition (Fig. 3). These findings indicate that the communication and interaction between distinct brain areas is enhanced under the influence of LSD, in line with the previous study of Tagliazucchi et al., where, amongst other findings, LSD was found to increase global integration by enhancing the level of communication between normally distinct brain networks^8^. Similar effects could be observed with psilocybin^56,60^. Interestingly, we also observed an increase in the basal integration in the music condition under LSD, while music during PCB condition led to a slight decrease in the basal integration. This opposing effect of music in the LSD versus PCB conditions could be related to an accentuated psychological response to music under psychedelics, as observed more generally in the psychedelic research literature^9,25,26^. The effect of music on brain activity in the placebo condition appeared to be more consistent with a generic ‘focused’ brain response – as suggested by a decrease in brain-wide integration and a narrowing of the repertoire of activity^61,62^. Music could be characterized as a type of (felt) intrinsic perturbation under LSD but perhaps less so under placebo, where it is more likely to be witnessed more as an external object. That there was less of a difference between the LSD and placebo condition in the final resting state scan (post-music), could be due to a waning effect of the drug (i.e. a pharmacokinetic factor) - as described in the Materials and Methods section, the third and final fMRI session (rest after music) was more temporally distanced to the subjective peak effect of the drug than the first two sessions -, or a residual effect of having just listened to music, e.g. stabilising mind and brain dynamics under LSD, such that they differ less from those of the placebo condition. It would be useful to test these speculations in the future with more experiments.

It was evident that almost every node revealed a marked difference in PILI values under LSD versus placebo (see Supplementary Tables S2 and S3) – and this was evident across all three scanning conditions (rest, rest with music, rest after music). A higher PILI value indicates that the perturbed node shows increased sensitivity and stronger reaction to a model perturbation and requires longer recovery time to return to normal baseline activity. This suggests there is a diminished ability of the brain to homeostatically ‘right itself’ after perturbation under LSD. It is well established that slowness of recovery to perturbation is a key property of critical systems, where it is sometimes referred to in the literature as “critical slowing”^63^. That the brain should exhibit critical slowing under psychedelics was recently hypothesised in a narrative review on the effects of psychedelics on global brain function (and note figure 2 in this article)^32^. The present findings therefore provide important empirical support for this principle.

Heightened sensitivity and the stronger reaction to a model perturbation in the LSD state models is also consistent with the work of Schartner et al.^36^, where elevated measures of MEG-recorded spontaneous or resting state brain complexity was found under psychedelics using an approach not unrelated to that of Massimini and colleagues^13,15,64,65^, who used TMS and complexity measures to characterise (diminished) states of consciousness. The here described effect of a simulated perturbation to LSD fMRI data could be regarded as a logical extension of these previous studies, where actual brain stimulation may be difficult to perform under a potent psychedelic. Moreover, the finding of elevated brain complexity is consistent with the finding of Schartner et al.^36^, Atasoy et al.^33^ as well as the entropic brain hypothesis^58,59^, which stipulates that within reasonable bounds, the complexity or entropy of spontaneous activity indexes the richness of conscious experience, where greater ‘richness’ implies greater diversity and depth.

Analyzing the perturbation-elicited differences on a local node and network level (Fig. 4 and Table 1), we found that some brain regions and networks were more dominant regarding differences in PILI than others. For example, the limbic network yielded the highest perturbation-elicited differences between the LSD and the PCB state models indicating an enhanced sensitivity of this network under the effect of LSD. Within this network, the cingulate cortex showed a remarkably large sensitivity (p < 10^−8^, effect size: 0.51 in music condition). The cingulate cortex, and the limbic system more generally, are both implicated in emotional processing^66^. Moreover, they are both also implicated in the brain action of psychedelics^67–70^. Interestingly, limbic brain regions, especially the medial temporal lobe, have been associated with producing transient dreamlike states with visual hallucinations, similar to psychedelic-like phenomena, upon electrical depth stimulation^71–74^, also supporting the involvement of these brain regions in psychedelic visions. The here presented finding of enhanced sensitivity to a model perturbation of the limbic network supports the well known effect of LSD to facilitate emotional arousal^75^. One could infer that heightened sensitivity of the limbic circuitry in particular is implicated in the heightened emotional responsivity that has been found in relation to psychedelic therapy^25,76,77^. The release of emotional content is thought to be a key aspect of the therapeutic action of psychedelic therapy^75,76^. Abnormal functioning of the limbic circuitry is well reported in mood disorders– and depression in particular^78,79^ which has been the target of psychedelic therapy^75,76^.

Two other networks, the visual network and the default mode network (DMN), were strongly altered by LSD, consistent with previous studies reporting changes in the functioning of visual areas and in the functional properties of the DMN under LSD^6,8^. Consistent with this result, brain changes involving visual regions have been found to correlate with eyes-closed imagery under LSD^6^, while changes in DMN properties have been found to correlate with high-level characteristics of the experience, including ego dissolution^8^.

Finally, in order to understand the level of variation across brain nodes in the perturbation response, we analyzed the perturbation response variability by looking at the variance over nodes of the perturbation-elicited responses. Larger variance over brain nodes means higher heterogeneity across brain regions. A larger response variability signifies that each brain region is becoming more independent in its activity after a strong model perturbation. We found that the response variability was significantly higher in all three scanning conditions under LSD than PCB (Fig. 6), which indicates an enhanced diversity in brain dynamics, as also previously suggested for the LSD state^33^. This effect is consistent with what one would expect from a breakdown in the usual hierarchical constraints governing global brain function. Interestingly, abnormal hierarchical organization has previously been associated with neuropathological disorders such as depression, with changes in multimodal network organization^80^ as well as psychosis and schizophrenia, with connectivity disturbances afflicting hierarchical brain organization^80^ leading to attenuated top-down cognitive control^81^. Furthermore autism also has been found to relate to differences in this multimodal network hierarchy^82^. The relationship between hierarchical organization in the brain and criticality (including critical slowing) was the focus of a recent major review on the acute and potential therapeutic action of psychedelics^32^ – and flattened functional hierarchy in the brain has recently been observed in formal ‘gradient-based’ analyses applied to the present dataset^83^.

The present study’s results suggest fundamental changes in brain dynamics and complexity under the influence of psychedelic drugs, consistent with the brain moving closer to a critical regime in which the brain is exquisitely sensitive to perturbation. These findings are therefore consistent with recent^32^ and older theoretical models of the effects of psychedelics on global brain function^59,84^. They also bear significant relevance to principles of psychedelic psychotherapy, where great emphasis is placed on the importance of context, or ‘set and setting’, as a principal modulator of outcomes^85^. More plainly, the present findings of increased brain sensitivity to perturbation under LSD could be interpreted as related to evidence-based assumptions^25^ about increased emotional sensitivity to environmental and other contextual factors (such as music) under psychedelics^85^.

The present version of the model allows us to understand how the global changes induced by LSD (i.e., global coupling) interact with the connectome and produce different network dynamics. The main limitation of the model is its homogeneity. In this model, all the brain regions were assumed to have the same intrinsic dynamics (a = 0). Therefore, within this model, the differences in the dynamics of the brain regions were a consequence of the different effective connectivity of the regions. The model could be extended by introducing heterogeneity in local dynamics (i.e., by allowing the parameter a to vary between brain regions, thus requiring the estimation of N new model parameters). This extension might be useful to investigate local changes produced by LSD. A further limitation of the model is its limited frequency range. Since the model was constructed based on BOLD signals, it can only produce slow frequencies. Probing the model with MEG signals could provide insights on how LSD affects the different frequency bands of brain activity.

In summary, by exploring the underlying mechanistic properties of the whole-brain dynamics in the LSD state using a novel in silico perturbational approach, we have provided important new insights into global brain function underlying a possible altered state of consciousness that could bear relevance to our understanding of brain function and conscious states more generally. Importantly, the perturbational approach based on whole-brain modelling allows for the exploration of characteristic changes in whole-brain dynamics in ways that are extremely challenging to do via in vivo experiments. Furthermore, the here presented results enrich our understanding of how psychedelic drugs may have therapeutic utility and suggest future research directions, in which the neural mechanisms underlying their clinical use can be further explored.

## Acknowledgements

BMJ, AP-A, AS and GD are supported by AWAKENING Using whole-brain models perturbational approaches for predicting external stimulation to force transitions between different brain states (ref. PID2019-105772GB-I00, AEI FEDER EU), funded by the Spanish Ministry of Science, Innovation and Universities (MCIU), State Research Agency (AEI) and European Regional Development Funds (FEDER), the HBP SGA3 Human Brain Project Specific Grant Agreement 3 (Grant Agreement No. 945539), funded by the EU H2020 FET Flagship program, the SGR Research Support Group support (ref. 2017 SGR 1545), funded by the Catalan Agency for Management of University and Research Grants (AGAUR), the SEMAINE ERA-Net NEURON Project, a Juan de la Cierva fellowship (IJCI-2014-21066) from the Spanish Ministry of Economy and Competitiveness, a Juan de la Cierva fellowship from the Spanish Ministry of Economy and Competitiveness (FPDI2013-17045), and by the FLAG-ERA JTC (PCI2018-092891).

RC-H is supported by the Alex Mosley Charitable Trust, Ad Astra Chandaria Foundation, Nikean Foundation, Tim Ferriss, Alexander and Bohdana Tamas and Anton Bilton. SA and MLK are supported by the ERC Consolidator Grant: CAREGIVING (n. 615539), Center for Music in the Brain, funded by the Danish National Research Foundation (DNRF117), and Centre for Eudaimonia and Human Flourishing funded by the Pettit and Carlsberg Foundations.

## Author contributions

BMJ and GD designed the study. BMJ, SA, AP-A and AS performed data analyses and numerical simulations. LR, MK and RC-H conducted fMRI experiments, pre-processed and provided the data. MLK provided the DTI data. BMJ wrote the first version of the manuscript. All authors contributed significantly to the writing of the article and agreed to the final version.

## Appendix A. Supplementary Information

