## Supplementary Information for "Increased sensitivity to strong perturbations in a whole-brain model of LSD"

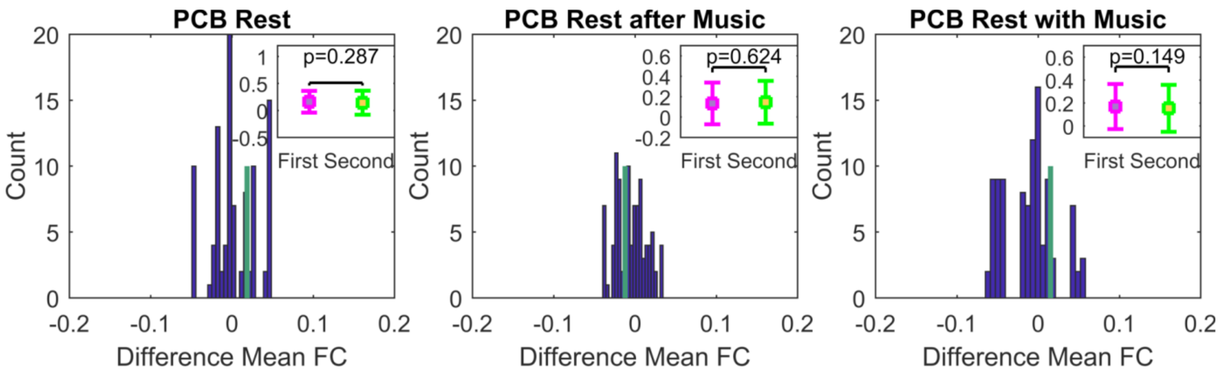


**Supplementary Figure S1: Differences Mean FCs Placebo - Division into first session PCB and second session PCB.** Here the differences between the group of participants who received PCB in their first session (group "First") vs. the group of participants who received PCB in their second session (group "Second") are analyzed. The differences of the mean of the upper triangle FC matrices (group average) between the groups "First" and "Second" are shown for each of the three conditions. The histograms (dark blue) represent the distribution of the test-statistic under the null-hypothesis of no difference between groups, whereas the green lines show the differences of the means of the empirical FC matrices. In the upper right corner of each panel the mean FCs and their standard deviations are shown. None of the differences are significantly different, meaning that the FC matrices present no differences between the two groups.

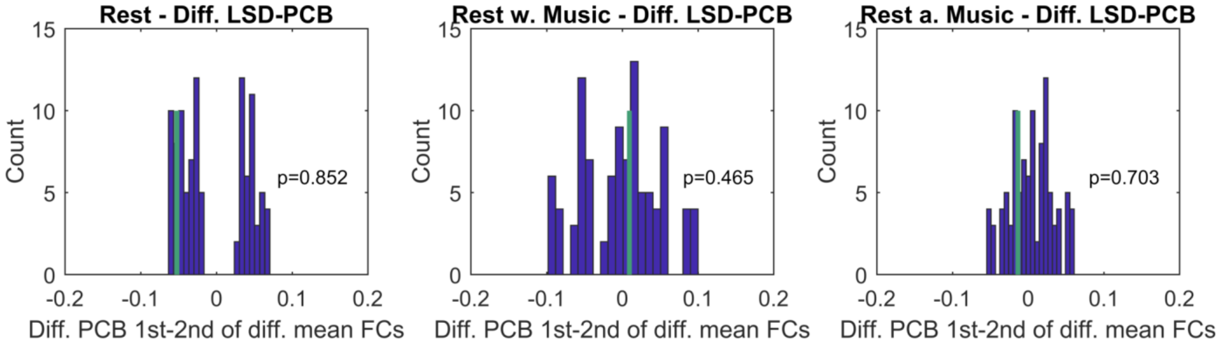


**Supplementary Figure S2: Differences Mean FCs LSD-PCB - Differences between first session PCB and second session PCB.** Here the differences between the group of participants who received PCB in their first session (group "First", here abbreviated with "1st") vs. the group of participants who received PCB in their second session (group "Second", here abbreviated with "2nd") are analyzed regarding the differences between the LSD and PCB groups. The participants were divided into two groups ("First" and "Second") and within those groups the differences of FC matrices between the LSD and PCB conditions were analyzed. Then, the mean FC differences between the two groups of these differences were calculated and tested for significance. The histograms (dark blue) represent the distribution of the test-statistic under the null-hypothesis of no difference between groups, meaning there is no difference between the groups "First" and "Second" regarding the differences between LSD and PCB. The green lines show the exact same differences of the empirical data. None of the differences are significantly different, meaning that there is no difference between the two groups regarding the differences between the LSD and PCB state.

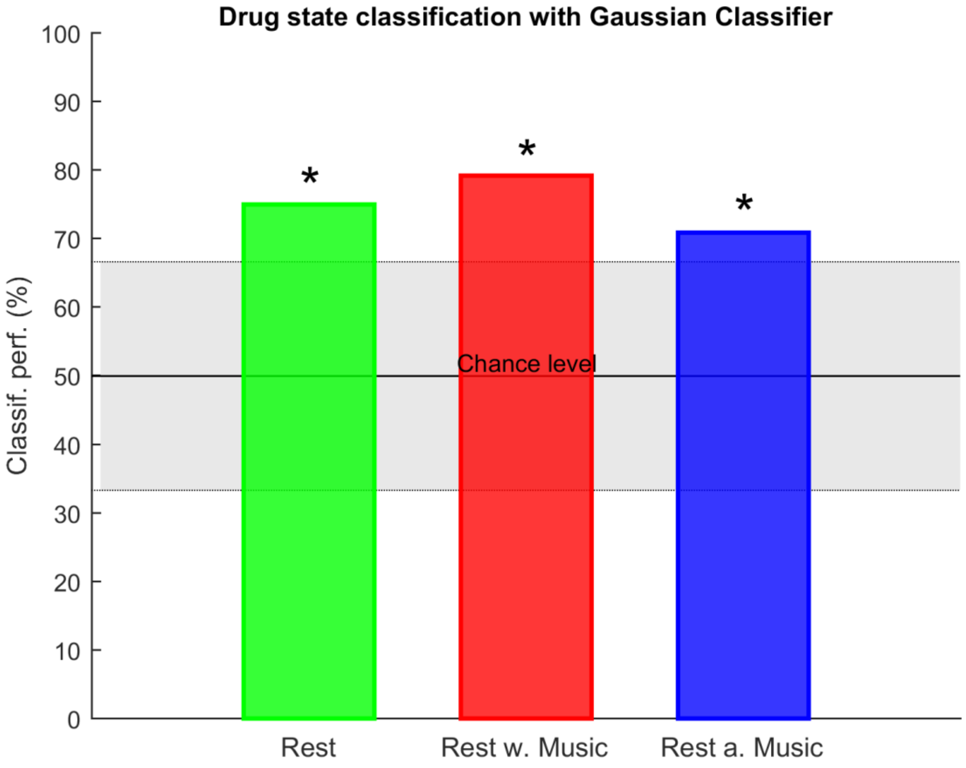


**Supplementary Figure S3: Drug state classification performance.** Here the classification performance using a Gaussian classifier is shown for all 3 scanning conditions (rest, rest with music and rest after music) (see *Methods*). The accuracy of the drug state (LSD or PCB) prediction exceeds in all 3 conditions the 95th percentile of chance level (75% for rest, 79,17% for rest with music and 70,83% for rest after music).

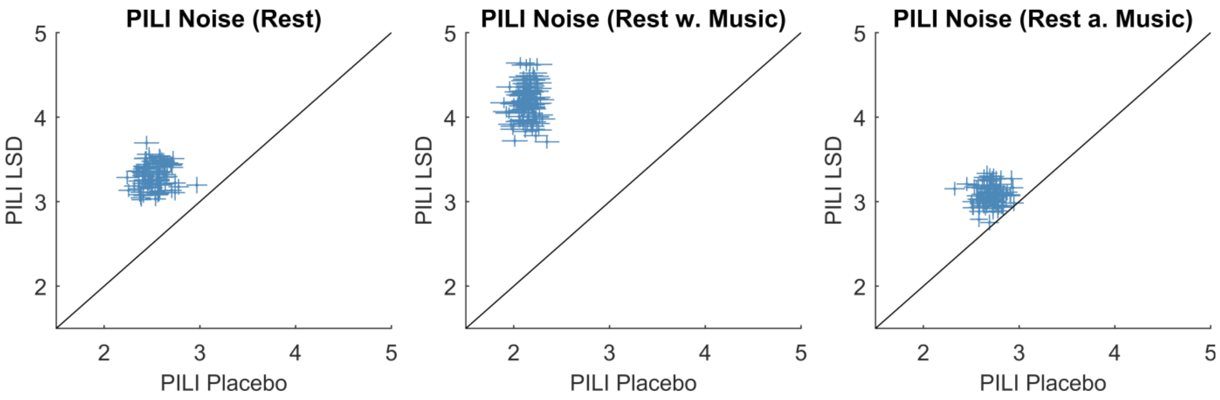


**Supplementary Figure S4: PILI - Node level analysis for noise protocol.** Here the mean and the standard error of the mean (SEM) of the PILI values over trials are shown for each of the three scanning conditions for the LSD and the PCB state for all 90 brain regions. The vertical error bars represent the SEM for the PCB state and horizontal error bars represent the errors for the LSD state. Also in the noise protocol, the global differences between the LSD and PCB induced brain states were amplified in the rest with music condition. Node-by-node analysis with corresponding p-values can be found in Table S2.

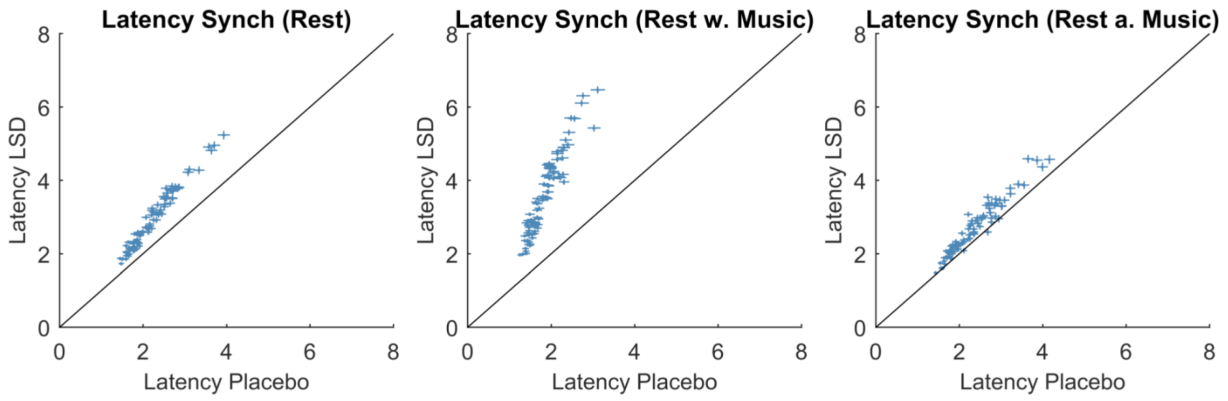


**Supplementary Figure S5: Latencies - Node level analysis for synchronization protocol.** Here the mean and the standard error of the mean (SEM) of the latency values over trials are shown for each of the three scanning conditions for the LSD and the PCB state for all 90 brain regions. The latencies indicate the time it takes for the perturbed signal to go back to the basal state. The vertical error bars represent the SEM for the PCB state and horizontal error bars represent the errors for the LSD state. In all 3 conditions higher latencies can be observed in the LSD state (Rest1: 88/90 nodes significantly higher latency in LSD, Music: 90/90 nodes significantly higher latency in LSD, Rest2: 62/90 nodes significantly higher latency in LSD).

As for the noise protocol, much less nodes show significantly higher latency: Rest1: 7/90 nodes significantly higher latency in LSD, Music: 34/90 nodes significantly higher latency in LSD, Rest2: 6/90 nodes significantly higher latency in LSD.

| AAL regions of interest | | | |
|---|---|---|---|
| **ROI Number** | **ROI name** | **cortical/subcortical** | **RSN** |
| 1 | 'Precentral_L' | cortical | 1,2,3,4,7 |
| 2 | 'Frontal_Sup_L' | cortical | 2,3,4,7 |
| 3 | 'Frontal_Sup_Orb_L' | cortical | 1,2,3,4 |
| 4 | 'Frontal_Mid_L' | cortical | 1,2,3,4,6,7 |
| 5 | 'Frontal_Mid_Orb_L' | cortical | 1,2,6 |
| 6 | 'Frontal_Inf_Oper_L' | cortical | 1,2,6 |
| 7 | 'Frontal_Inf_Tri_L' | cortical | 1,2,3,4 |
| 8 | 'Frontal_Inf_Orb_L' | cortical | 1,2,3,4 |
| 9 | 'Rolandic_Oper_L' | cortical | 1,2,6 |
| 10 | 'Supp_Motor_Area_L' | cortical | 1,2,6 |
| 11 | 'Olfactory_L' | cortical | 1,2,3,4 |
| 12 | 'Frontal_Sup_Medial_L' | cortical | 1,2,3,4 |
| 13 | 'MedialOFC_L' | cortical | 1,2,4 |
| 14 | 'Rectus_L' | cortical | 1,2,3,4 |
| 15 | 'Insula_L' | cortical | 1,2,4,6 |
| 16 | 'Cingulum_Ant_L' | cortical | 1,2,6 |
| 17 | 'Cingulum_Mid_L' | cortical | 4,7 |
| 18 | 'Cingulum_Post_L' | cortical | 4,7 |
| 19 | 'Hippocampus_L' | subcortical | 1,2,4,7 |
| 20 | 'ParaHippocampal_L' | subcortical | 1,2,4,7 |
| 21 | 'Amygdala_L' | subcortical | 6 |
| 22 | 'Calcarine_L' | cortical | 6 |
| 23 | 'Cuneus_L' | cortical | 1,2,4,6 |
| 24 | 'Lingual_L' | cortical | 1,2 |
| 25 | 'Occipital_Sup_L' | cortical | 1,6 |
| 26 | 'Occipital_Mid_L' | cortical | 1,6 |
| 27 | 'Occipital_Inf_L' | cortical | 1,6 |
| 28 | 'Fusiform_L' | cortical | 1,6 |
| 29 | 'Postcentral_L' | cortical | 1,2,4,6,7 |
| 30 | 'Parietal_Sup_L' | cortical | 1,2,4,6,7 |
| 31 | 'Parietal_Inf_L' | cortical | 1,2,4,6 |
| 32 | 'SupraMarginal_L' | cortical | 1,2,4 |
| 33 | 'Angular_L' | cortical | 1,2,4,7 |
| 34 | 'Precuneus_L' | cortical | 1,2,4,7 |
| 35 | 'Paracentral_Lobule_L' | cortical | 1,2 |
| 36 | 'Caudate_L' | subcortical | 1,2 |
| 37 | 'Putamen_L' | subcortical | 1,6 |
| 38 | 'Pallidum_L' | subcortical | 0 |
| 39 | 'Thalamus_L' | subcortical | 1,5,6 |
| 40 | 'Heschl_L' | cortical | 1,5,6 |
| 41 | 'Temporal_Sup_L' | cortical | 6 |
| 42 | 'Temporal_Pole_Sup_L' | cortical | 1,2,3,4,5,6,7 |
| 43 | 'Temporal_Mid_L' | cortical | 1,2,3,4,5,6,7 |
| 44 | 'Temporal_Pole_Mid_L' | cortical | 1,5 |
| 45 | 'Temporal_Inf_L' | cortical | 1,2,3,5 |
| 46 | 'Temporal_Inf_R' | cortical | 1,2,3,5 |
| 47 | 'Temporal_Pole_Mid_R' | cortical | 1,5 |
| 48 | 'Temporal_Mid_R' | cortical | 1,5 |

| | | | |
|---|---|---|---|
| 49 | 'Temporal_Pole_Sup_R' | cortical | 1,2,3,5 |
| 50 | 'Temporal_Sup_R' | cortical | 2,3,5 |
| 51 | 'Heschl_R' | cortical | 1,2,3,5 |
| 52 | 'Thalamus_R' | subcortical | 1,3,5 |
| 53 | 'Pallidum_R' | subcortical | 3,5 |
| 54 | 'Putamen_R' | subcortical | 5 |
| 55 | 'Caudate_R' | subcortical | 1,3,5,6 |
| 56 | 'Paracentral_Lobule_R' | cortical | 1,3,5,6 |
| 57 | 'Precuneus_R' | cortical | 3,4,7 |
| 58 | 'Angular_R' | cortical | 3,7 |
| 59 | 'SupraMarginal_R' | cortical | 2,3,5,7 |
| 60 | 'Parietal_Inf_R' | cortical | 3,5,7 |
| 61 | 'Parietal_Sup_R' | cortical | 1,2,3,4,7 |
| 62 | 'Postcentral_R' | cortical | 1,2,3,4,7 |
| 63 | 'Fusiform_R' | cortical | 1,2,3,4,7 |
| 64 | 'Occipital_Inf_R' | cortical | 2,3,4,7 |
| 65 | 'Occipital_Mid_R' | cortical | 1,2,3 |
| 66 | 'Occipital_Sup_R' | cortical | 1,2,3 |
| 67 | 'Lingual_R' | cortical | 1,2,3,4,5,7 |
| 68 | 'Cuneus_R' | cortical | 1,2,3,4,5,7 |
| 69 | 'Calcarine_R' | cortical | 7 |
| 70 | 'Amygdala_R' | subcortical | 4 |
| 71 | 'ParaHippocampal_R' | subcortical | 0 |
| 72 | 'Hippocampus_R' | subcortical | 0 |
| 73 | 'Cingulum_Post_R' | cortical | 0 |
| 74 | 'Cingulum_Mid_R' | cortical | 0 |
| 75 | 'Cingulum_Ant_R' | cortical | 0 |
| 76 | 'Insula_R' | cortical | 0 |
| 77 | 'Rectus_R' | cortical | 0 |
| 78 | 'MedialOFC_R' | cortical | 0 |
| 79 | 'Frontal_Sup_Medial_R' | cortical | 7 |
| 80 | 'Olfactory_R' | cortical | 4,7 |
| 81 | 'Supp_Motor_Area_R' | cortical | 1,4,6,7 |
| 82 | 'Rolandic_Oper_R' | cortical | 1,2,4,7 |
| 83 | 'Frontal_Inf_Orb_R' | cortical | 1,4,6,7 |
| 84 | 'Frontal_Inf_Tri_R' | cortical | 1,4,6,7 |
| 85 | 'Frontal_Inf_Oper_R' | cortical | 1,2,3,4,5,7 |
| 86 | 'Frontal_Mid_Orb_R' | cortical | 1,2,3,4,5,7 |
| 87 | 'Frontal_Mid_R' | cortical | 1,6 |
| 88 | 'Frontal_Sup_Orb_R' | cortical | 1,6 |
| 89 | 'Frontal_Sup_R' | cortical | 1,2,3,5,6 |
| 90 | 'Precentral_R' | cortical | 1,2,3,5,6 |

**Supplementary Table S1: List of AAL regions of interest**. Complete list of ROI regions with information about whether the region is cortical or subcortical. Furthermore the association of each ROI to the corresponding RSNs is described (1: Default mode network, 2: Executive control network, 3: Dorsal attention network, 4: Ventral attention network, 5: Visual network, 6: Limbic network, 7:

Somatomotor network, 0: Corpus Callosum and other subcortical structures (not considered in the RSN analysis)).

| Rest | | | Rest with Music | | | Rest after Music | | |
|---|---|---|---|---|---|---|---|---|
| **Brain region** | **p-value** | **Effect size** | **Brain region** | **p-value** | **Effect size** | **Brain region** | **p-value** | **Effect size** |
| Cuneus R | 5.68e-26* | 0.1924 | Putamen R | 8.76e-127* | 0.4373 | Frontal Inf Oper R | 3.28e-08* | 0.1009 |
| Occipital Sup L | 1.58e-25* | 0.1907 | Caudate R | 2.26e-125* | 0.4348 | Temporal Sup L | 3.94e-08* | 0.1003 |
| Rectus R | 2.61e-25* | 0.1898 | Calcarine R | 1.80e-124* | 0.4332 | Amygdala L | 4.12e-08* | 0.1002 |
| Frontal Sup R | 7.96e-25* | 0.1878 | Supp Motor Area L | 3.14e-120* | 0.4257 | Frontal Sup L | 4.44e-08* | 0.0999 |
| Pallidum L | 1.57e-24* | 0.1866 | Occipital Sup L | 1.11e-119* | 0.4247 | Amygdala R | 5.17e-08* | 0.0994 |
| Frontal Sup Orb R | 7.94e-24* | 0.1837 | Paracentral Lobule L | 1.12e-118* | 0.4229 | Frontal Mid R | 8.06e-08* | 0.0980 |
| Frontal Mid Orb R | 8.77e-24* | 0.1836 | Temporal Pole Sup L | 1.90e-118* | 0.4224 | Occipital Inf L | 8.30e-08* | 0.0979 |
| MedialOFC L | 1.42e-23* | 0.1827 | Hippocampus R | 4.81e-118* | 0.4217 | Frontal Sup Medial L | 8.31e-08* | 0.0979 |
| Caudate L | 3.40e-23* | 0.1811 | MedialOFC L | 1.51e-117* | 0.4208 | Cingulum Post L | 1.81e-07* | 0.0953 |
| Cingulum Mid R | 3.97e-23* | 0.1808 | Amygdala R | 4.55e-116* | 0.4181 | MedialOFC R | 1.84e-07* | 0.0952 |
| Insula L | 4.35e-22* | 0.1764 | Putamen L | 3.46e-110* | 0.4072 | Parietal Sup R | 2.05e-07* | 0.0948 |
| Parietal Sup R | 7.27e-22* | 0.1755 | Frontal Mid Orb L | 1.26e-109* | 0.4061 | ParaHippocampal L | 2.23e-07* | 0.0946 |
| Parietal Sup L | 5.42e-21* | 0.1716 | Frontal Inf Orb L | 3.36e-108* | 0.4034 | Occipital Sup R | 2.97e-07* | 0.0936 |
| Amygdala L | 9.79e-21* | 0.1705 | Lingual R | 2.20e-107* | 0.4019 | Occipital Sup L | 3.36e-07* | 0.0932 |
| Hippocampus L | 1.02e-20* | 0.1704 | Insula R | 5.82e-105* | 0.3972 | Caudate L | 3.65e-07* | 0.0929 |
| Occipital Inf L | 1.56e-20* | 0.1696 | Temporal Pole Sup R | 9.94e-105* | 0.3968 | Supp Motor Area L | 4.39e-07* | 0.0922 |
| Occipital Mid L | 1.81e-20* | 0.1693 | Fusiform L | 1.12e-104* | 0.3967 | Olfactory R | 1.08e-06* | 0.0890 |
| Paracentral Lobule L | 2.71e-20* | 0.1685 | Occipital Mid L | 1.82e-103* | 0.3944 | Pallidum L | 1.44e-06* | 0.0880 |
| Frontal Sup L | 3.55e-20* | 0.1680 | Olfactory L | 1.00e-96* | 0.3810 | Calcarine L | 1.49e-06* | 0.0879 |
| Frontal Mid Orb L | 4.84e-20* | 0.1674 | Frontal Inf Tri L | 5.33e-95* | 0.3775 | Olfactory L | 1.67e-06* | 0.0874 |
| Frontal Inf Orb L | 6.60e-20* | 0.1668 | Occipital Mid R | 5.40e-92* | 0.3714 | Paracentral Lobule R | 3.01e-06* | 0.0853 |
| Frontal Inf Orb R | 2.66e-19* | 0.1640 | Insula L | 1.96e-91* | 0.3702 | Temporal Pole Sup R | 3.32e-06* | 0.0849 |
| Fusiform R | 2.69e-19* | 0.1640 | Thalamus L | 7.82e-91* | 0.3690 | MedialOFC L | 6.16e-06* | 0.0825 |
| Frontal Sup Orb L | 2.81e-19* | 0.1639 | Temporal Sup R | 1.28e-90* | 0.3686 | Frontal Sup Orb L | 6.74e-06* | 0.0822 |

| | | | | | | | | |
|---|---|---|---|---|---|---|---|---|
| Frontal Sup Medial R | 8.70e-19* | 0.1616 | Frontal Inf Oper R | 7.43e-88* | 0.3628 | Parietal Sup L | 8.32e-06* | 0.0814 |
| Cuneus L | 1.62e-18* | 0.1603 | Cuneus L | 5.53e-87* | 0.3609 | Cingulum Post R | 1.27e-05* | 0.0797 |
| Lingual R | 3.43e-18* | 0.1588 | Fusiform R | 4.08e-86* | 0.3591 | Temporal Mid L | 2.87e-05* | 0.0764 |
| MedialOFC R | 7.24e-18* | 0.1572 | Frontal Mid R | 7.93e-86* | 0.3585 | Temporal Sup R | 3.46e-05* | 0.0756 |
| Temporal Inf R | 1.71e-17* | 0.1554 | ParaHippocampal R | 9.09e-86* | 0.3583 | Occipital Mid R | 4.49e-05* | 0.0745 |
| Insula R | 2.57e-17* | 0.1545 | Occipital Sup R | 3.39e-84* | 0.3550 | Caudate R | 5.66e-05* | 0.0735 |
| Supp Motor Area L | 2.82e-16* | 0.1494 | Calcarine L | 6.62e-84* | 0.3543 | Thalamus L | 7.07e-05* | 0.0726 |
| Pallidum R | 2.85e-16* | 0.1493 | Temporal Mid R | 5.23e-83* | 0.3524 | Putamen R | 7.50e-05* | 0.0723 |
| ParaHippocampal L | 1.19e-15* | 0.1462 | Temporal Pole Mid L | 5.46e-83* | 0.3524 | SupraMarginal R | 9.26e-05* | 0.0714 |
| Temporal Mid R | 1.52e-15* | 0.1456 | Hippocampus L | 6.03e-83* | 0.3523 | Fusiform L | 1.10e-04* | 0.0706 |
| Temporal Mid L | 1.55e-15* | 0.1456 | Occipital Inf L | 1.08e-82* | 0.3517 | Precentral L | 1.15e-04* | 0.0704 |
| Occipital Inf R | 1.92e-15* | 0.1451 | Cuneus R | 1.69e-82* | 0.3513 | Insula R | 1.27e-04* | 0.0700 |
| Occipital Mid R | 8.10e-15* | 0.1418 | Frontal Inf Tri R | 2.96e-82* | 0.3508 | Postcentral L | 1.46e-04* | 0.0693 |
| Precentral R | 8.86e-15* | 0.1416 | Thalamus R | 3.19e-82* | 0.3507 | Lingual L | 1.92e-04* | 0.0681 |
| Caudate R | 9.01e-15* | 0.1415 | Lingual L | 4.21e-81* | 0.3482 | Paracentral Lobule L | 2.46e-04* | 0.0669 |
| Frontal Mid L | 9.25e-15* | 0.1415 | Parietal Sup L | 5.66e-81* | 0.3479 | Heschl L | 3.12e-04* | 0.0658 |
| Temporal Pole Sup R | 3.26e-14* | 0.1385 | Temporal Mid L | 1.00e-80* | 0.3474 | Rectus R | 3.47e-04* | 0.0653 |
| Paracentral Lobule R | 5.42e-14* | 0.1373 | Rolandic Oper R | 7.34e-79* | 0.3433 | Insula L | 4.10e-04* | 0.0645 |
| Frontal Inf Oper L | 5.63e-14* | 0.1372 | Precentral R | 1.10e-77* | 0.3406 | Frontal Sup Orb R | 4.26e-04* | 0.0643 |
| Frontal Mid R | 9.49e-14* | 0.1360 | Paracentral Lobule R | 3.43e-77* | 0.3395 | Temporal Pole Mid L | 5.10e-04* | 0.0635 |
| Amygdala R | 1.08e-13* | 0.1357 | Frontal Mid Orb R | 6.08e-77* | 0.3390 | Frontal Inf Orb R | 6.21e-04 | 0.0625 |
| Parietal Inf R | 2.14e-13* | 0.1340 | Frontal Inf Oper L | 2.93e-72* | 0.3282 | Postcentral R | 9.31e-04 | 0.0604 |
| Rolandic Oper L | 2.97e-13* | 0.1332 | Temporal Inf L | 2.08e-69* | 0.3215 | Temporal Inf R | 9.51e-04 | 0.0603 |
| Precentral L | 4.10e-13* | 0.1324 | Postcentral L | 1.18e-65* | 0.3125 | Putamen L | 1.30e-03 | 0.0587 |
| Fusiform L | 4.91e-13* | 0.1320 | Rolandic Oper L | 2.71e-65* | 0.3116 | Frontal Inf Orb L | 1.30e-03 | 0.0587 |
| Rolandic Oper R | 7.26e-13* | 0.1310 | Precentral L | 6.35e-65* | 0.3107 | Frontal Inf Oper L | 4.40e-03 | 0.0520 |

| | | | | | | | | |
|---|---|---|---|---|---|---|---|---|
| Frontal Inf Oper R | 7.83e-13* | 0.1308 | Temporal Pole Mid R | 1.53e-63* | 0.3072 | SupraMarginal L | 4.53e-03 | 0.0518 |
| Temporal Sup L | 1.24e-12* | 0.1296 | Frontal Mid L | 5.59e-62* | 0.3033 | Frontal Mid Orb R | 4.89e-03 | 0.0514 |
| Postcentral R | 2.69e-12* | 0.1277 | Temporal Inf R | 1.16e-61* | 0.3025 | Cuneus L | 0.012 | 0.0461 |
| Heschl L | 7.00e-12* | 0.1252 | Amygdala L | 3.58e-61* | 0.3013 | Temporal Mid R | 0.022 | 0.0420 |
| Temporal Pole Sup L | 1.46e-11* | 0.1233 | Pallidum R | 6.18e-61* | 0.3007 | Frontal Mid L | 0.023 | 0.0416 |
| Temporal Pole Mid R | 3.44e-11* | 0.1210 | ParaHippocampal L | 1.17e-60* | 0.3000 | Rolandic Oper L | 0.052 | 0.0356 |
| Frontal Inf Tri L | 3.67e-11* | 0.1208 | Temporal Sup L | 1.64e-60* | 0.2996 | Lingual R | 0.093 | 0.0306 |
| ParaHippocampal R | 1.09e-10* | 0.1178 | Parietal Inf L | 4.87e-55* | 0.2853 | Temporal Pole Mid R | 0.096 | 0.0304 |
| Parietal Inf L | 1.20e-10* | 0.1176 | Frontal Inf Orb R | 9.74e-54* | 0.2818 | Angular L | 0.110 | 0.0292 |
| Heschl R | 1.42e-09* | 0.1105 | Heschl R | 3.00e-53* | 0.2804 | Parietal Inf L | 0.177 | 0.0246 |
| SupraMarginal L | 2.65e-09* | 0.1087 | Angular L | 6.76e-53* | 0.2795 | Rolandic Oper R | 0.192 | 0.0238 |
| Temporal Sup R | 3.78e-09* | 0.1076 | Pallidum L | 1.50e-47* | 0.2645 | Heschl R | 0.210 | 0.0229 |
| Temporal Inf L | 7.90e-09* | 0.1054 | Occipital Inf R | 6.20e-47* | 0.2627 | Occipital Inf R | 0.237 | 0.0216 |
| Hippocampus R | 1.43e-08* | 0.1035 | Angular R | 2.88e-44* | 0.2548 | Frontal Inf Tri R | 0.237 | 0.0216 |
| SupraMarginal R | 1.44e-07* | 0.0960 | SupraMarginal R | 2.60e-42* | 0.2489 | Cuneus R | 0.250 | 0.0210 |
| Frontal Inf Tri R | 3.89e-07* | 0.0926 | Postcentral R | 3.34e-37* | 0.2327 | Frontal Inf Tri L | 0.359 | 0.0168 |
| Postcentral L | 3.48e-06* | 0.0847 | Parietal Sup R | 2.10e-36* | 0.2301 | Parietal Inf R | 0.543 | 0.0111 |
| Temporal Pole Mid L | 1.43e-05* | 0.0792 | SupraMarginal L | 7.92e-35* | 0.2248 | Pallidum R | 0.585 | 0.0100 |
| Angular L | 1.58e-04* | 0.0690 | Heschl L | 1.28e-25* | 0.1910 | Thalamus R | 0.618 | 0.0091 |
| Angular R | 4.71e-04* | 0.0638 | Parietal Inf R | 8.35e-20* | 0.1663 | Angular R | 0.698 | 0.0071 |

* statistically significant after Bonferroni correction

**Supplementary Table S2: Node level PILI differences - synchronization protocol.** In this table brain nodes are ordered for each condition by p-values - from smallest to largest -, based on the PILI differences for the synchronization protocol between LSD and PCB by perturbing each specific node at a time. This table represents a continuation of Table 1, where here the regions 21 up to 90 are shown ordered by the size of their p-value with their corresponding effect sizes.

| Rest | | | Rest with Music | | | Rest after Music | | |
|---|---|---|---|---|---|---|---|---|
| **Brain region** | **p-value** | **Effect size** | **Brain region** | **p-value** | **Effect size** | **Brain region** | **p-value** | **Effect size** |
| Frontal Sup Orb R | 1.14e-07* | 0.0968 | Occipital Mid L | 1.99e-23* | 0.1821 | Rectus R | 5.07e-05* | 0.0740 |
| Rolandic Oper R | 9.38e-07* | 0.0895 | Cuneus R | 6.62e-21* | 0.1712 | Frontal Mid Orb L | 5.56e-05* | 0.0736 |
| Frontal Sup Medial R | 1.85e-06* | 0.0871 | Frontal Mid R | 4.53e-20* | 0.1675 | Frontal Sup Medial L | 7.03e-05* | 0.0726 |
| Frontal Sup Orb L | 3.05e-06* | 0.0852 | Olfactory L | 1.29e-19* | 0.1654 | Rolandic Oper R | 1.69e-04* | 0.0687 |
| Temporal Sup L | 5.84e-06* | 0.0827 | Caudate L | 4.97e-19* | 0.1627 | SupraMarginal L | 4.20e-04* | 0.0644 |
| Olfactory L | 7.00e-06* | 0.0820 | Supp Motor Area R | 1.79e-18* | 0.1601 | Insula L | 8.28e-04 | 0.0610 |
| Hippocampus L | 7.36e-06* | 0.0818 | Rolandic Oper L | 3.44e-18* | 0.1588 | Cingulum Mid L | 9.26e-04 | 0.0605 |
| Cingulum Ant L | 8.29e-06* | 0.0814 | Thalamus L | 1.13e-17* | 0.1563 | Cingulum Ant R | 1.25e-03 | 0.0589 |

| | | | | | | | | |
|---|---|---|---|---|---|---|---|---|
| Calcarine L | 1.26e-05* | 0.0797 | MedialOFC R | 2.40e-17* | 0.1547 | Caudate R | 1.43e-03 | 0.0582 |
| Amygdala L | 1.37e-05* | 0.0794 | Temporal Mid R | 2.42e-17* | 0.1547 | Cuneus R | 1.74e-03 | 0.0572 |
| MedialOFC L | 1.41e-05* | 0.0793 | Parietal Inf L | 4.33e-17* | 0.1534 | Cingulum Post L | 2.13e-03 | 0.0561 |
| Putamen R | 2.64e-05* | 0.0767 | Insula L | 5.80e-17* | 0.1528 | Occipital Mid L | 3.46e-03 | 0.0534 |
| Occipital Sup R | 2.88e-05* | 0.0764 | Hippocampus L | 1.53e-16* | 0.1507 | Precuneus R | 4.82e-03 | 0.0515 |
| Temporal Pole Mid L | 4.33e-05* | 0.0747 | Frontal Sup Medial L | 2.90e-16* | 0.1493 | Occipital Sup R | 6.10e-03 | 0.0501 |
| Frontal Mid R | 4.41e-05* | 0.0746 | Precuneus L | 9.74e-16* | 0.1466 | Lingual L | 8.39e-03 | 0.0481 |
| Paracentral Lobule R | 5.38e-05* | 0.0737 | Calcarine L | 1.87e-15* | 0.1451 | Thalamus L | 0.011 | 0.0464 |
| Pallidum R | 5.80e-05* | 0.0734 | Frontal Sup Orb L | 1.92e-15* | 0.1451 | Frontal Inf Orb R | 0.013 | 0.0456 |
| Rectus L | 6.63e-05* | 0.0728 | Putamen R | 4.06e-15* | 0.1434 | Heschl R | 0.019 | 0.0430 |
| Rolandic Oper L | 7.14e-05* | 0.0725 | Frontal Sup R | 4.39e-15* | 0.1432 | Angular L | 0.019 | 0.0427 |
| Thalamus R | 7.24e-05* | 0.0725 | Frontal Inf Orb L | 4.49e-15* | 0.1431 | Heschl L | 0.021 | 0.0420 |
| Olfactory R | 7.92e-05* | 0.0721 | Putamen L | 7.36e-15* | 0.1420 | Frontal Inf Tri R | 0.023 | 0.0416 |
| Angular L | 1.10e-04* | 0.0706 | Rectus R | 9.12e-15* | 0.1415 | Frontal Sup Orb L | 0.032 | 0.0393 |
| Occipital Inf R | 1.14e-04* | 0.0705 | Cingulum Post R | 1.04e-14* | 0.1412 | Paracentral Lobule L | 0.034 | 0.0388 |
| Cingulum Post R | 1.25e-04* | 0.0700 | Rectus L | 1.20e-14* | 0.1409 | Fusiform L | 0.039 | 0.0378 |
| Temporal Mid L | 1.42e-04* | 0.0695 | Frontal Inf Orb R | 1.95e-14* | 0.1397 | Occipital Sup L | 0.041 | 0.0373 |
| Frontal Mid Orb R | 1.43e-04* | 0.0694 | Frontal Mid Orb R | 2.59e-14* | 0.1391 | Temporal Pole Mid L | 0.042 | 0.0371 |
| Parietal Sup R | 2.66e-04* | 0.0666 | Angular L | 2.86e-14* | 0.1388 | MedialOFC L | 0.043 | 0.0370 |
| Caudate L | 4.01e-04* | 0.0646 | MedialOFC L | 3.20e-14* | 0.1386 | Calcarine R | 0.052 | 0.0355 |
| Supp Motor Area L | 4.31e-04* | 0.0643 | Precentral R | 5.22e-14* | 0.1374 | Cingulum Mid R | 0.053 | 0.0354 |
| Hippocampus R | 4.72e-04* | 0.0638 | SupraMarginal L | 6.48e-14* | 0.1369 | ParaHippocampal R | 0.064 | 0.0339 |
| Occipital Mid L | 8.33e-04 | 0.0610 | Supp Motor Area L | 1.20e-13* | 0.1354 | Rolandic Oper L | 0.067 | 0.0335 |
| Fusiform L | 9.39e-04 | 0.0604 | Frontal Sup L | 2.41e-13* | 0.1337 | ParaHippocampal L | 0.073 | 0.0328 |
| Angular R | 1.58e-03 | 0.0577 | Cuneus L | 3.05e-13* | 0.1331 | MedialOFC R | 0.078 | 0.0322 |
| Precentral L | 1.72e-03 | 0.0572 | Rolandic Oper R | 3.57e-13* | 0.1327 | Supp Motor Area L | 0.079 | 0.0321 |

| | | | | | | | | |
|---|---|---|---|---|---|---|---|---|
| Cingulum Mid R | 1.75e-03 | 0.0571 | Temporal Inf R | 3.60e-13* | 0.1327 | Frontal Sup R | 0.093 | 0.0307 |
| Parietal Inf L | 1.97e-03 | 0.0565 | Occipital Sup L | 3.79e-13* | 0.1326 | Hippocampus R | 0.095 | 0.0305 |
| Occipital Mid R | 2.12e-03 | 0.0561 | ParaHippocampal L | 4.79e-13* | 0.1320 | Temporal Sup L | 0.096 | 0.0304 |
| Temporal Pole Mid R | 2.37e-03 | 0.0555 | Postcentral R | 5.28e-13* | 0.1318 | Pallidum R | 0.096 | 0.0304 |
| Frontal Inf Tri L | 2.40e-03 | 0.0554 | Temporal Pole Mid R | 6.56e-13* | 0.1312 | Frontal Mid L | 0.102 | 0.0299 |
| Insula R | 2.51e-03 | 0.0552 | Temporal Mid L | 1.64e-12* | 0.1289 | Frontal Inf Oper R | 0.105 | 0.0296 |
| Frontal Mid Orb L | 2.76e-03 | 0.0547 | Fusiform L | 2.28e-12* | 0.1281 | Angular R | 0.113 | 0.0289 |
| Temporal Pole Sup R | 2.87e-03 | 0.0544 | Occipital Sup R | 2.75e-12* | 0.1276 | Amygdala R | 0.120 | 0.0284 |
| ParaHippocampal R | 2.88e-03 | 0.0544 | Insula R | 2.92e-12* | 0.1275 | Olfactory R | 0.136 | 0.0272 |
| Calcarine R | 3.48e-03 | 0.0534 | Pallidum R | 5.96e-12* | 0.1256 | Precentral R | 0.140 | 0.0270 |
| Paracentral Lobule L | 3.54e-03 | 0.0533 | Occipital Inf L | 6.39e-12* | 0.1254 | Temporal Sup R | 0.149 | 0.0264 |
| Cuneus R | 3.76e-03 | 0.0529 | Precuneus R | 7.05e-12* | 0.1252 | Frontal Mid Orb R | 0.162 | 0.0255 |
| Temporal Pole Sup L | 3.90e-03 | 0.0527 | Paracentral Lobule L | 9.12e-12* | 0.1245 | Paracentral Lobule R | 0.173 | 0.0249 |
| Frontal Inf Tri R | 4.39e-03 | 0.0520 | Cingulum Post L | 9.35e-12* | 0.1244 | Amygdala L | 0.181 | 0.0244 |
| Temporal Mid R | 4.40e-03 | 0.0520 | Olfactory R | 1.48e-11* | 0.1232 | Temporal Mid R | 0.187 | 0.0241 |
| Pallidum L | 4.49e-03 | 0.0519 | Parietal Sup L | 3.03e-11* | 0.1213 | Cingulum Post R | 0.193 | 0.0238 |
| Frontal Sup Medial L | 4.54e-03 | 0.0518 | Fusiform R | 3.03e-11* | 0.1213 | Lingual R | 0.201 | 0.0234 |
| Cingulum Ant R | 4.74e-03 | 0.0516 | Postcentral L | 3.46e-11* | 0.1210 | Occipital Mid R | 0.214 | 0.0227 |
| Parietal Inf R | 5.09e-03 | 0.0511 | Frontal Sup Medial R | 9.00e-11* | 0.1184 | Frontal Sup L | 0.222 | 0.0223 |
| Heschl R | 6.40e-03 | 0.0498 | Occipital Mid R | 1.36e-10* | 0.1172 | Caudate L | 0.252 | 0.0209 |
| Insula L | 6.47e-03 | 0.0497 | Temporal Pole Sup R | 3.48e-10* | 0.1146 | Putamen R | 0.261 | 0.0205 |
| Thalamus L | 7.71e-03 | 0.0486 | Temporal Sup R | 4.73e-10* | 0.1137 | Temporal Pole Sup L | 0.286 | 0.0195 |
| Frontal Inf Oper R | 7.74e-03 | 0.0486 | Angular R | 6.50e-10* | 0.1128 | Temporal Inf R | 0.354 | 0.0169 |
| Heschl L | 7.75e-03 | 0.0486 | Frontal Inf Oper R | 8.63e-10* | 0.1120 | Frontal Sup Medial R | 0.367 | 0.0165 |
| Temporal Inf L | 8.00e-03 | 0.0484 | Caudate R | 1.08e-09* | 0.1113 | Temporal Inf L | 0.371 | 0.0163 |
| Fusiform R | 0.014 | 0.0448 | Cingulum Ant L | 1.37e-09* | 0.1106 | Parietal Inf R | 0.395 | 0.0155 |

| | | | | | | | | |
|---|---|---|---|---|---|---|---|---|
| SupraMarginal L | 0.016 | 0.0441 | Cingulum Mid R | 1.45e-09* | 0.1105 | Occipital Inf R | 0.417 | 0.0148 |
| Precuneus L | 0.022 | 0.0419 | Pallidum L | 1.47e-09* | 0.1104 | Frontal Inf Tri L | 0.425 | 0.0146 |
| Caudate R | 0.023 | 0.0414 | Parietal Inf R | 1.76e-09* | 0.1099 | Temporal Pole Mid R | 0.439 | 0.0141 |
| Temporal Inf R | 0.024 | 0.0413 | Frontal Sup Orb R | 1.84e-09* | 0.1097 | Olfactory L | 0.441 | 0.0141 |
| Postcentral R | 0.029 | 0.0398 | Frontal Inf Oper L | 3.73e-09* | 0.1076 | Thalamus R | 0.448 | 0.0139 |
| Parietal Sup L | 0.031 | 0.0393 | Frontal Mid Orb L | 4.09e-09* | 0.1074 | Parietal Inf L | 0.480 | 0.0129 |
| SupraMarginal R | 0.032 | 0.0392 | ParaHippocampal R | 7.54e-09* | 0.1055 | Temporal Mid L | 0.488 | 0.0127 |
| Cingulum Post L | 0.035 | 0.0385 | Temporal Inf L | 7.70e-09* | 0.1054 | Rectus L | 0.514 | 0.0119 |
| ParaHippocampal L | 0.035 | 0.0384 | Lingual L | 9.42e-09* | 0.1048 | Precuneus L | 0.551 | 0.0109 |
| Precentral R | 0.040 | 0.0376 | Frontal Inf Tri L | 9.77e-09* | 0.1047 | Supp Motor Area R | 0.560 | 0.0106 |
| Postcentral L | 0.043 | 0.0968 | Precentral L | 1.24e-08* | 0.1821 | Postcentral R | 0.588 | 0.0740 |
| Frontal Sup R | 0.045 | 0.0895 | Calcarine R | 1.72e-08* | 0.1712 | Occipital Inf L | 0.592 | 0.0736 |
| Cuneus L | 0.047 | 0.0871 | Frontal Mid L | 1.95e-08* | 0.1675 | Cuneus L | 0.600 | 0.0726 |
| Cingulum Mid L | 0.052 | 0.0852 | Hippocampus R | 3.32e-08* | 0.1654 | Hippocampus L | 0.655 | 0.0687 |
| Supp Motor Area R | 0.056 | 0.0827 | Heschl L | 4.13e-08* | 0.1627 | Cingulum Ant L | 0.658 | 0.0644 |
| Amygdala R | 0.067 | 0.0820 | Thalamus R | 5.12e-08* | 0.1601 | Fusiform R | 0.687 | 0.0610 |
| Lingual L | 0.071 | 0.0818 | Occipital Inf R | 8.98e-08* | 0.1588 | Parietal Sup L | 0.705 | 0.0605 |
| Precuneus R | 0.094 | 0.0814 | Temporal Sup L | 9.16e-08* | 0.1563 | Calcarine L | 0.734 | 0.0589 |
| Lingual R | 0.096 | 0.0797 | Frontal Inf Tri R | 1.40e-07* | 0.1547 | Frontal Mid R | 0.738 | 0.0582 |
| Rectus R | 0.099 | 0.0794 | SupraMarginal R | 2.08e-07* | 0.1547 | Frontal Inf Orb L | 0.753 | 0.0572 |
| MedialOFC R | 0.118 | 0.0793 | Temporal Pole Sup L | 2.30e-07* | 0.1534 | Parietal Sup R | 0.769 | 0.0561 |
| Frontal Inf Oper L | 0.126 | 0.0767 | Parietal Sup R | 2.42e-07* | 0.1528 | Precentral L | 0.799 | 0.0534 |
| Occipital Sup L | 0.157 | 0.0764 | Amygdala L | 2.84e-07* | 0.1507 | Postcentral L | 0.819 | 0.0515 |
| Frontal Inf Orb L | 0.296 | 0.0747 | Paracentral Lobule R | 3.90e-07* | 0.1493 | Frontal Sup Orb R | 0.823 | 0.0501 |
| Occipital Inf L | 0.344 | 0.0746 | Amygdala R | 5.60e-07* | 0.1466 | Pallidum L | 0.861 | 0.0481 |
| Frontal Inf Orb R | 0.392 | 0.0737 | Heschl R | 2.49e-06* | 0.1451 | SupraMarginal R | 0.867 | 0.0464 |

| | | | | | | | | |
|---|---|---|---|---|---|---|---|---|
| Temporal Sup R | 0.511 | 0.0734 | Lingual R | 3.96e-06* | 0.1451 | Insula R | 0.871 | 0.0456 |
| Frontal Sup L | 0.556 | 0.0728 | Cingulum Ant R | 8.99e-06* | 0.1434 | Temporal Pole Sup R | 0.887 | 0.0430 |
| Frontal Mid L | 0.587 | 0.0725 | Temporal Pole Mid L | 1.54e-05* | 0.1432 | Frontal Inf Oper L | 0.995 | 0.0427 |
| Putamen L | 0.669 | 0.0725 | Cingulum Mid L | 4.39e-03 | 0.1431 | Putamen L | 0.997 | 0.0420 |

* statistically significant after Bonferroni correction

**Supplementary Table S3: Node level PILI differences - noise protocol.** In this table brain nodes are ordered for each condition by p-values - from smallest to largest -, based on the PILI differences for the noise protocol between LSD and PCB by perturbing each specific node at a time. All 90 brain regions are shown in order by the size of their p-value with their corresponding effect sizes.

| Differences PILI RSNs | | | |
|---|---|---|---|
| | Rest | Rest with Music | Rest after Music |
| RSN | p-value | p-value | p-value |
| Default mode network | 2.98e-08* | 2.42e-21* | 0.002* |
| Executive control network | 3.80e-06* | 8.49e-17* | 0.010 |
| Dorsal attention network | 3.85e-05* | 3.02e-12* | 0.012 |
| Ventral attention network | 9.55e-06* | 1.64e-13* | 0.020 |
| Visual network | 1.55e-04* | 2.29e-09* | 0.030 |
| Limbic network | 4.83e-06* | 6.51e-12* | 0.004* |
| Somatomotor network | 9.67e-06* | 2.58e-11* | 0.023 |

* statistically significant after Bonferroni correction

**Supplementary Table S4: RSN level PILI differences.** In this table the p-values for each condition are shown on an RSN level based on the PILI differences for the synchronization protocol between LSD and PCB. All networks show statistically significant differences, except for the Rest after Music condition, where 5 out of 7 networks don't survive the Bonferroni correction for multiple comparisons.
